# Lenacapavir prevents production of infectious HIV-1 by abrogating immature virus assembly

**DOI:** 10.64898/2026.03.02.709178

**Authors:** Clifton L Ricaña, Savannah G Brancato, Carolyn M Highland, Fatemeh Ekbataniamiri, Krisztina Ambrus, Juan S Rey, Mia Faerch, Bruce E Torbett, Juan R Perilla, Robert A Dick

## Abstract

The HIV-1 capsid effector Lenacapavir (LEN) acts by disrupting the early (reverse transcription and nuclear entry) and late (assembly and maturation) stages of the viral lifecycle. The early stage requires an intact Capsid consisting of a lattice of capsid protein (CA) hexamers and pentamers. Phenylalanine-Glycine (FG) containing nuclear pore proteins interact with Capsid hexamers at their FG pockets, facilitating nuclear entry. Disruption of Capsid lattice stability, or competitive binding to the FG pocket, by LEN blocks infection. Here, we provide insight into the effects of LEN on the late stage. Using a combination of cryo-EM structure determination, *in vitro* assembly, and *in situ* viral assays, we determined that treatment of producer cells with LEN abrogates the production of infectious virus via multiple mechanisms. Previous studies have shown that HIV-1 produced from LEN treated cells have improperly formed Capsids. However, how LEN interacts with the CA domain of the Gag polyprotein during assembly, prior to maturation, is unclear. Using a viral protease (PR) defective HIV-1 clone, which traps the Gag lattice in an immature state, we found that LEN dramatically remodulates the CA domain. The resulting CA layer was mature-like despite the absence of PR cleavage. We dubbed this unnatural state as “premature” lattice. Proper immature and mature lattice formation requires inositol hexakisphosphate (IP6), but our cryo-EM work revealed a lack of IP6 in the premature lattice. These findings provide insight into the mechanisms by which LEN prevents infectious HIV-1 production.

## Introduction

HIV-1 assembly, budding, and maturation are highly regulated processes (reviewed (1-3)). During assembly, the structural protein Gag interacts with the viral genomic RNA (vgRNA), the cellular metabolite inositol hexakisphosphate (IP6), and the inner leaflet of the cellular plasma membrane (PM) resulting in the formation of a lattice of Gag hexamers. Ultimately, this results in the release of an immature virus. Subsequently, the viral protease (PR) cleaves Gag into its constituent domains, restructuring the protein lattice. The liberated capsid (CA) domain assembles into a different hexameric lattice, with 12 distinct pentamers in the shape of a fullerene cone (Capsid) that encapsulates the vgRNA and enzymatic proteins, converting the virus to an infectious mature form. After release into a target cell (reviewed (4)), Capsid is trafficked through the cytoplasm. In the Capsid, vgRNA is reverse transcribed into DNA and eventually released from Capsid for subsequent integration into the cell genome (reviewed (5, 6)).

A current prevailing model is that Capsid docks to nuclear pores (reviewed (7)) via interactions with various nucleoporins (Nups) (8). Most Nups have regions of unstructured peptide motifs comprised of Phenylalanine-Glycine (FG) repeats which form a hydrogel allowing for directed movement through the nuclear envelope. The CA_NTD_ of mature Capsid hexamers has an FG binding pocket which interacts with the Nups (8). This pocket is a target of Capsid effectors including PF-3450074 (PF74) (9, 10) and the first-in-class FDA approved compound, Lenacapavir (LEN, marketed by Gilead Sciences as Sunlenca and Yeztugo) (11, 12).

LEN binding to FG pockets of Capsid hexamers can block interactions with FG containing proteins such as Nup153 and CPSF6, thus preventing nuclear entry (11, 12). Binding of LEN can also alter Capsid stability leading to incomplete reverse transcription (13-18). While the effects of LEN on the mature Capsid core are well documented, less is known about the effects of LEN on immature assembly, budding, and maturation. Leveraging advances in cryo-EM structure determination of retroviral lattices (19-29), we investigated the effects of LEN on the lattice of *in vitro* assembled immature HIV-1 virus like particles (VLPs) and *in situ* VLPs budded from cells. We report here that LEN binds to and disrupts assembly of immature virus resulting in the formation of an aberrant lattice that fails to properly employ IP6 or correctly mature. These structural effects are sufficient to account for the reported loss of infectivity.

## Results

### The effect of LEN on CASPNC immature lattice assembly

The development of HIV-1 Capsid binding effectors has heightened the importance of understanding the molecular interactions between effectors and the mature Capsid lattice. Since the CA protein that comprises Capsid is also an integral domain of Gag, and thus part of the immature lattice, understanding Gag interactions with effectors is equally important (Fig. S1A-B). For this reason, we tested the effect of LEN on immature lattice when added at various steps to our standard *in vitro* assembly (23) of purified CASPNC protein. This construct was chosen as it contains the main structured region of the immature Gag lattice. When monomeric CASPNC (50 μM) was incubated with LEN (50 μM) before assembly, the VLPs were less complete compared to the 0 μM LEN control (DMSO) (Fig. S1C). When LEN was added during or after assembly, the assembled VLPs were morphologically indistinguishable from DMSO (Fig. S1C).

IP6 is a metabolite with cellular concentrations ranging from 10-40 μM (30) and is known to play a critical role in the formation of both the immature and mature HIV-1 lattices, as previously observed for both *in vitro*-assembled VLPs as well as *in situ*-formed virus (23, 31, 32). Many groups have described an interplay between IP6 and LEN on mature Capsid stability (13, 15, 33-35). As IP6 is also critical for immature lattice formation, we assessed the effect of adding LEN to immature CASPNC VLP assembly reactions with decreasing concentrations of IP6. At our established assembly concentration of IP6 (10 μM), addition of LEN did not visibly alter either the efficiency or gross morphology of the immature VLPs (Fig. S1D). However, at low IP6 concentrations and with the addition of LEN, we saw mature CASPNC assembled into tubular structures (Fig. S1D), consistent with previously described tubular assemblies of CA (13, 23).

### SPA structure determination of in vitro assembled HIV-1 immature lattice

Using SPA to interrogate *in vitro*-assembled HIV-1 immature CASPNC VLPs, we determined the structure of the hexameric lattice to a resolution of 1.9 Å (Fig. 1A, Fig. S2A-B, and Table S1), which allowed for unambiguous modeling of all side chains. The addition of DMSO did not alter particle morphology or structure compared to previously published works (20, 21, 29). Next, we determined the structure of *in vitro*-assembled immature VLPs in complex with LEN, achieving a global resolution of 2.0 Å (Fig. 1B, Fig. S2C-F, and Table S1). The VLPs in both the absence and presence of LEN displayed the same gross morphology and 2D classes (Fig. 1A-B and Fig. S2A-F), indicating that LEN did not visually disrupt normal lattice assembly. The final map contained extra density at the N-terminal domain (NTD) (Fig. 1B). Force-field parameterization of LEN (Fig. S3) and subsequent modeling into our cryo-EM map demonstrated that the extra density is consistent with LEN binding to the hydrophilic pocket formed by alpha helices (αH) αH3-5 and αH7 (Fig. 1B and Fig. S1A-B and S4) (11, 12, 36). In agreement with the previous report from Xiong and colleagues (29), we found that LEN remodels the CA_NTD_ but does not affect the CA_CTD_SP domains (Fig. 1D-E). Further analysis of the CA_NTD_ revealed that while there was considerable deviation of αH1 and αH3-6, αH2 remained relatively fixed (Fig. 1E). As αH2 comprises a three-fold interface, we analyzed the quaternary structure comprising six trimer interfaces of the immature lattice. From this more wholistic view, we observed that LEN binding causes a rotation along the trimer interface at αH2 (Fig. 1F and Movie S1).

**Figure 1.**
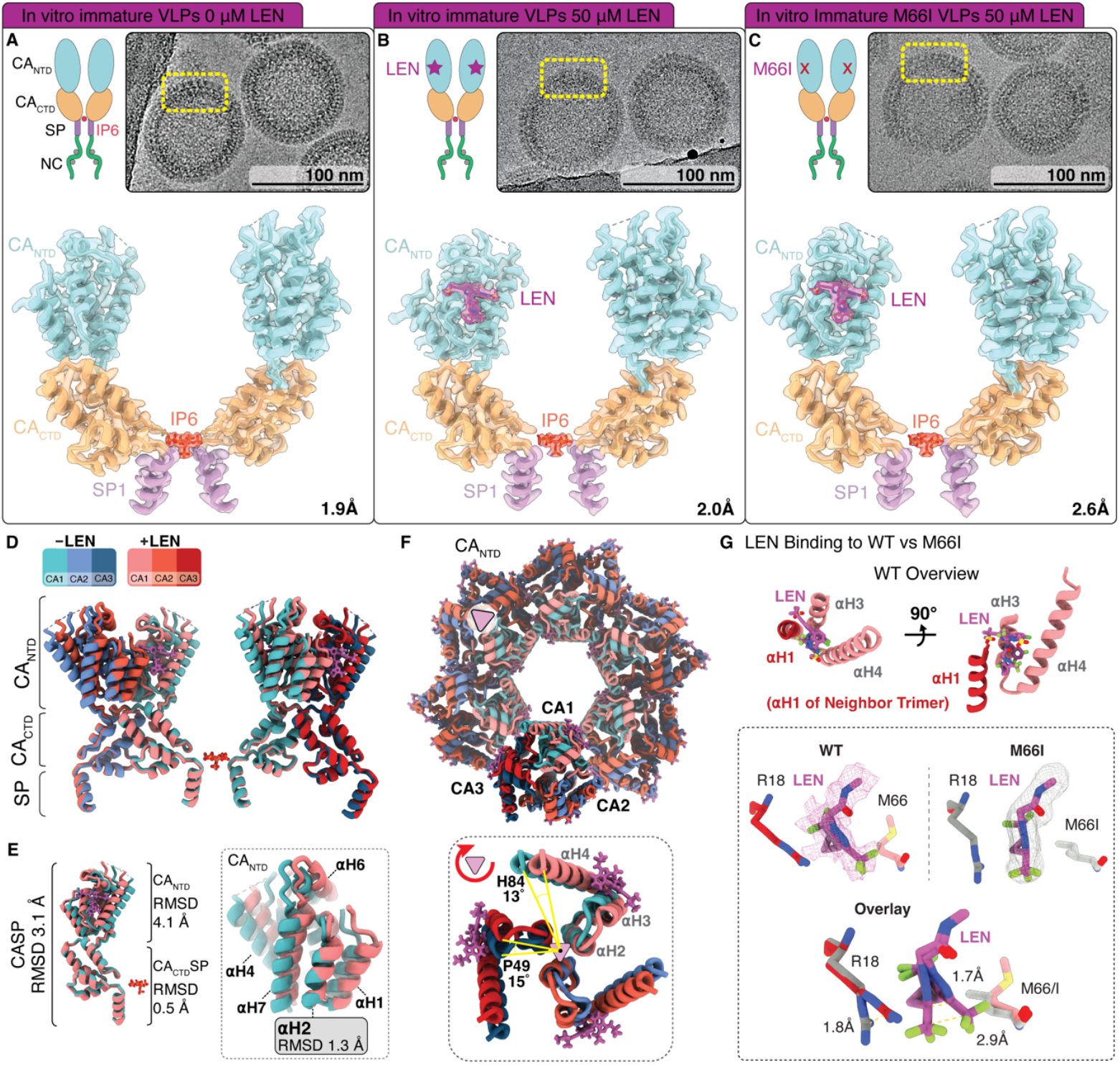
HIV-1 protein lattice analysis of in vitro assembled immature (CASPNC) particles. **A-C)** Top, left: Schematic of the structured regions of two monomers of the CASP hexamer (NC is unstructured). Top, right: Representative micrograph cutouts of assembled CASPNC particles. Bottom: Side view representation of the determined maps and models of two monomers of the immature hexamer. **A)** WT treatment with 0 μM LEN. **B)** WT treatment with 50 μM LEN. **C)** M66I treatment with 50 μM LEN. **D)** Side view of the four monomers comprising two opposing CTD-dimer interfaces. Note the alpha helices displaced by the addition of LEN at the NTD compared to the lack of displacement at the CTD-SP. **E)** Left: Side view of one monomer showing RMSD values at the different domains of CASP. Right: Zoom-in of the NTD showing alpha helical (αH2 to a lesser extent) displacement by LEN. **F)** Top: Top-down view of the NTDs of the central immature hexamer and neighboring monomers, together which comprise six three-fold interfaces. Bottom: Zoom-in of the NTDs showing rotation around αH2. Note clockwise rotation along the central axis of three αH2 helices. **G)** Top: Key residues involved in LEN binding (WT) with αH3 and αH4 of one trimer and αH1 of the neighboring trimer. Bottom: Binding of the M3a (pentalene-like) moiety of LEN to CASPNC WT compared to M66I. Note ability of the M3a moiety to rotate as well as the corresponding movement of the neighboring monomer R18 residue.

We modeled putative waters interacting with LEN and proximal side chains based on their bond angle and distance (Fig. S4A). We found that LEN binding causes significant displacement of water from the previously described hydrophilic pocket (36). Importantly, there are two regions where water is retained via interactions with amino acids and LEN. The first region is at a sulfonyl moiety (M2). The second is at the pentalene-like moiety (M3a) and the 3,5-difluorobenzene moiety (M3b) (Fig. S4B-E). Previous crystallography work showed that M3a coordinates a water with Q67, K70, and N183 (PDB: 6V2F) (11). Interestingly, our immature lattice structure shows that both halves of M3 coordinate a water with L56, V59, and additional waters (Fig. S4B, water #1). LEN also disrupts a water network involving R18 of αH1, resulting in a pi-stacking interaction between R18 and M3a. Taken together, our results show that LEN binding alters the hydration state of the protein and induces the movement of αH1 toward αH4 of the neighboring monomer (Fig S4A and C), resulting in the observed rotation of the CA_NTD_ around the αH2 threefold axis (Fig. 1F).

Understanding the molecular mechanisms by which resistance-associated mutations (RAMs) disrupt effector binding is important for the design of second-generation effectors. For LEN, one such RAM is M66I, which displays >230-fold reduced binding compared to WT CA (37). Using our *in vitro* system, we were able to saturate M66I assemblies to overcome the high K_D_ reported for the mutation and determined the structure of LEN bound to the immature M66I lattice to 2.6 Å (Fig. 1C, Fig. S2G-H, and Table S1). M66 interacts with the M3 moiety of LEN (Fig. S4B and D). Overlay of the WT and M66I cryo-EM maps and atomic models showed that the M3a moiety can exhibit a level of rotational freedom around the linking carbon (Fig. 1G). This rotational movement is further supported by the corresponding movement of the pi-bonded R18. In the mature lattice, the CTD of the adjacent monomer likely restricts rotation of M3a, leading to steric hindrance as previously described (37). This is the first reported structure of LEN bound to a lattice with a RAM.

### SPA structure determination of the in situ HIV-1 immature and mature lattices

To validate the *in vitro* assembled model, we first applied SPA to immature VLPs produced *in situ* (Fig. S5) in the presence of DMSO as a control, leading to a 2.3 Å structure (Fig. 2A, Fig. S5A-B, and Table S1). Briefly, a full-length, LTR-driven NL4.3 HIV^ΔEnv^ provirus with a catalytic site mutation in the viral protease (PR-D25A) (19) was used for production of these authentic immature VLPs (Fig. S6A). The VLPs from transfected DMSO-treated cells were purified and western blot analysis verified a lack of Gag processing (Fig. S6B-C). VLP gross morphology and determined structure show that the VLPs have an immature Gag lattice matching pervious reports (19-22) (Fig. 2A, Fig. S5A-B).

**Figure 2.**
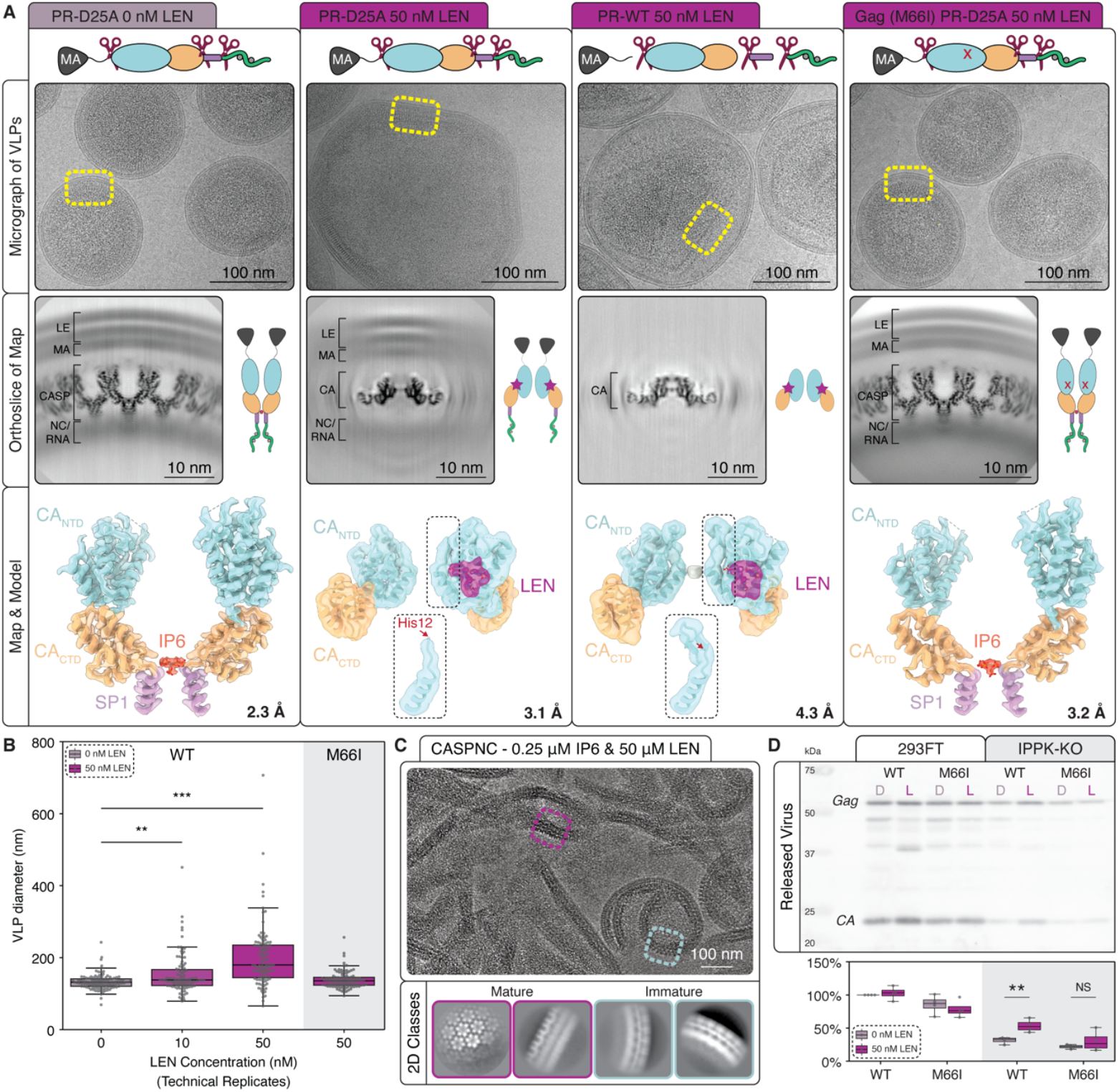
HIV-1 protein lattice analysis of in situ immature (PR-D25A) VLPs. **A)** Schematic of Gag cleavage sites as well as representative micrograph cutouts, orthoslices from the determined cryo-EM maps (LE lipid envelope), and atomic models laid within cryo-EM maps. Red arrow indicates His12 location. **B)** Quantification of VLP diameter across the horizontal axis (technical replicates, n = 105; Kruksal-Wallis Test, ** P<0.01, *** P<0.001). **C)** Representative cryo-EM micrograph and 2D classes from CASPNC *in vitro* assemblies using 0.25 μM IP6 and 50 μM LEN. Note 2D-classes consistent with mature lattice conformation in purple. **D)** Viral protein release from 293FT or 293FT-IPPK-KO cells pretreated for 24 h with DMSO (D) or 50 nM LEN (L) (lysate data in Fig. S8E). Release was assayed from concentrated spent media at 48 h post-transfection with HIV-1 provirus and no VSVg (n=4, Student’s T-Test with Welch’s correction, NS = Not significant, ** P<0.01).

Next, we determined the structure of *in situ* PR-D25A VLPs produced from cells pretreated with LEN (Fig. 2A, Fig. S5C-H, and Table S1). Compared to DMSO, VLPs from LEN pretreated cells were enlarged by two to three times, consistent with a previous report (38). These VLPs had a flatter protein lattice corresponding to the CA layer of Gag (Fig. 2A-B). At 10 nM LEN, VLPs appeared heterogeneous, and 2D classification showed regions of both immature lattice and unnatural mature-like lattice. Refinement of the immature and mature-like classes resulted in 2.7 Å and 4.8 Å maps, respectively (Fig. S5C-F). Strikingly, at 50 nM LEN, the VLPs displayed only mature-like lattice, resulting in a 3.1 Å map (Fig. 2A and Fig. S5G-H).

Interestingly, despite observing a mature-like lattice, there were no centralized cores normally observed following maturation. Additionally, the protein density was in close proximity to the viral membrane and did not have the β-turn at residues Pro1-His12, which is a hallmark of cleavage of CA from MA (Fig. 2A and Fig. S1A). Western blot analysis also confirmed a lack of cleavage (Fig. S6C). Importantly, we observed extra densities corresponding to LEN between the NTDs and CTDs of the hexameric lattice, consistent with previous reports of where LEN binds to the mature lattice (Fig. 2A) (11, 12, 26). These VLPs showed less curvature compared to DMSO controls, as depicted in the micrographs and by the orthoslices of the determined structure (Fig. 2A). Of note, in this sample with LEN, there was a lack of density corresponding to IP6 (Fig. 2A) which is consistent with the lack of observed pentamers (efficient assembly of which requires IP6) in the same sample and in our previously reported *in vitro* structure (26). As this lattice presents both immature and mature characteristics, we term this aberrant mature-like lattice as *premature*.

As a control, we determined the structure of *in situ* mature (PR-WT) VLPs from LEN pretreated cells to 4.3 Å (Fig. 2A, Fig. S7, and Table S1). The same Env-deficient provirus was used, but without the protease inactivating PR-D25A mutation (Fig. S6A). Similarly to premature VLPs, these mature VLPs were two to three times larger than those produced in the presence of DMSO, consistent with a previous report (38), and displayed flatter facets (Fig. S8A). The Capsid cores had a larger, more polyhedral morphology compared to DMSO (Fig. S8A). Western blot analysis and 2D classes confirmed mature lattice morphology, and the density detail is consistent with the FSC curves (Fig. S6C far right and Fig. S7A-B). Additionally, orthoslices of the determined structure demonstrate a lack of density corresponding to viral membrane indicating that the mature lattice has been cleaved from Gag as opposed to the premature lattice (Fig. 2A). The orthoslices further demonstrate that the lattice is flatter in the presence of LEN (Fig. 2A and Fig. S8A).

We determined the hexamer structure (3.7 Å) of *in vitro* assembled and LEN-bound Capsid-like particles (CLPs) (Fig. S7C-E and Table S1). These assemblies are very similar in gross morphology to authentic LEN-free Capsid cores, albeit less intact (Fig. S8B). The atomic structure with LEN modeled from the cryo-EM map of these CLPs readily docks into our 4.2 Å LEN-bound *in situ* cryo-EM map (Fig. 2A). Comparison of the atomic models between premature and mature lattices showed minimal deviation (RMSD 0.985 Å) (Fig S8C), and both these models are consistent with previously described mature lattices with LEN bound (11, 12, 26). Additionally, the CA_NTD_ of the premature lattice also shows minimal deviation from the *in vitro* derived immature lattice (RMSD 0.766 Å) (Fig. S8D). While the premature and immature NTDs display minimal deviation from each other, the bound LEN molecule does deviate at the M3a moiety, recapitulating the rotational freedom that was observed in the immature M66I LEN-bound model (Fig. 1G). To further verify that the observed premature lattice is specific to LEN binding to the NTD, we tested the PR-D25A M66I RAM. Even at 50 nM LEN, only the immature lattice morphology was observed (Fig. 2A and Fig. S5I-J). Our results demonstrate that *in situ* and in the absence of RAMs, LEN alters the HIV-1 assembly pathway to promote formation of an aberrant premature lattice.

### In vitro and in situ immature lattices in the presence of LEN are different

Our *in vitro* IP6 assembled CASPNC VLPs with LEN have an immature CA arrangement, while our *in situ* Gag (MACASPNCp6, i.e. full-length) VLPs released from cells pretreated with LEN have a mature-like CA arrangement defined above as premature. We predict that the primary reason for this observation is the difference in available IP6 *in vitro* compared to *in situ* during VLP assembly. Consistent with this hypothesis, there is a lack of strong density for IP6 at the center of the hexamers observed in the *in situ* premature and mature lattices in the presence of LEN (Fig. 2A). Normally, density corresponding to IP6 binding at the mature CA hexamer is located above and below the first turn of αH1, interacting with R18 and R18/K25 respectively (Fig. S7E) (26, 32). The density seen in Fig. 2A (PR-WT 50 nM LEN) is instead located in-plane with the first turn of αH1 and is not present in the off-symmetry axis hexamers. It is, therefore, likely an artifact of data processing. As noted above, the addition of LEN to *in vitro* immature assemblies with low IP6 results in tubular assemblies of CASPNC (Fig. S1D). We collected a small cryo-EM data set of this assembly and determined that these assemblies have a mature-like CA lattice as shown in the 2D classes (Fig. 2C).

We and others have shown that blocking the production of IP6 in cells via an IPMK- or IPPK-KO results in a ∼20-fold reduction in virus release (23, 39-41) Using the established IPPK-KO system, we measured VLPs release from cells treated with LEN. When treated with 50 nM LEN, a moderate recovery of release from IPPK-KO cells was observed (Fig. 2D and Fig. S8E). In summary, our results suggest that LEN competes with or bypasses IP6-induced assembly in cells, resulting in the observed formation of the aberrant premature lattice.

### LEN packaged into virus during assembly blocks subsequent infection

To understand the consequences of LEN induced aberrant immature assembly and maturation for entry of HIV-1 into target cells, we measured infectivity using flow cytometry. Producer cells were pretreated with increasing concentrations of LEN and then transfected with a pseudotyped VSVg HIV^ΔEnv^ provirus containing an EGFP reporter (Fig. S6A and S9A). Similar pretreatment infectivity assays have shown that LEN is inhibitory at picomolar concentrations (11, 12). We aimed to reduce the possibility of LEN carry-over from pretreated cells/media to those being infected. Virus-containing medium harvested from transfected producer cells was purified to removed LEN via pelleting through a 20% sucrose cushion and subsequent washing with PBS (Fig. 3A and Fig. S9A). Half of the washed virus was used to infect effector-naïve cells, half of which were scored for GFP fluorescence as a proxy for infection (Fig. S9B). Pretreatment of producing cells with increasing concentrations of LEN resulted in a corresponding decrease in infectivity of the released virus (Fig. 3B). As a control, virus produced from cells pretreated with 0 nM LEN (DMSO) was used to infect target cells that had been treated with 5 nM LEN 24 h prior to infection. This converse experiment resulted in a similar loss in infectivity, compared to the loss in infectivity when virus was produced in cells in the presence of the effector (Fig. 3B).

**Figure 3.**
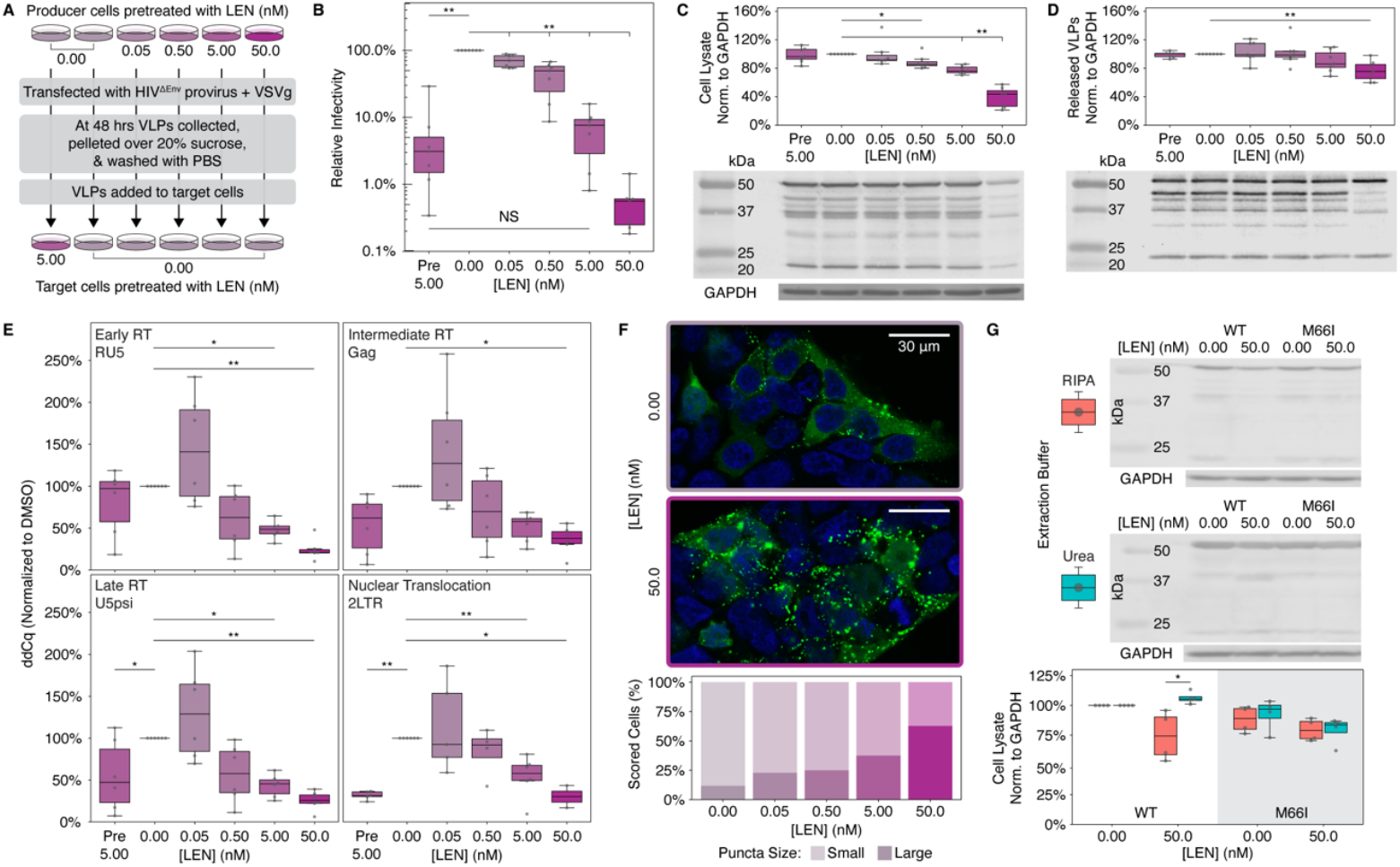
LEN packaged into VLPs during assembly in cells is sufficient for the observed downstream inhibitory effects. **A)** Abridged depiction of the experimental design (full diagram in Fig. S9B). VLPs were collected at 48 h post-transfection from cells pretreated with increasing LEN concentrations for 24 h. VLPs were pelleted through a 20% sucrose cushion and washed with PBS before downstream infection of LEN-treated or DMSO-treated cells. **B)** Infectivity as measured by the percentage of green-fluorescent target cells detected by flow cytometry; n = 7; Kruksal-Wallis Test; ** *P*<0.01). **C)** Total αCA signal from producer cell lysate. Top: Quantification (n = 7; Kruksal-Wallis Test; * *P*<0.05, ** *P*<0.01). Bottom: Representative western blot. **D)** Total αCA signal from purified virus released from cells in **C**. Top: Quantification (n = 7; Kruksal-Wallis Test; ** *P*<0.01). Bottom: Representative western blot. **E)** Relative copy number (ddCt, normalized to DMSO-treatment) for early RT (RU5, collected 6 h post-infection), intermediate RT (Gag, collected at 48 h post-infection), late RT (U5psi, collected at 48 h post-infection), and nuclear translocation (2LTR circles, collected at 48 h post-infection) of cells infected with virus from **D** (n = 4-6; Kruksal-Wallis Test; * *P*<0.05, ** *P*<0.01). **F)** Top: Representative confocal images of 24 h DMSO or LEN pretreated cells that were transfected with a Gag-mNeonGreen fusion protein (imaged 24 h post-transfection). Bottom: Quantification of the percentage of cells with either small or large puncta (n > 100). **G)** Lysate from producer cells pretreated with 50 nM LEN for 24 h (collected 48 h post-transfection with HIV-1 provirus without VSVg). Top: Representative western blot of lysate extracted with RIPA lysis buffer. Middle: Representative western blot of lysate extracted with 8 M Urea lysis buffer. Bottom: Quantification of lysate extracted with either RIPA or 8 M Urea lysis buffer (n = 4; Kruksal-Wallis Test; * *P*<0.05).

To determine if the observed reduction in infection was due to a reduction in protein expression, release, or reverse transcriptase (RT) activity, we employed several methods. To address protein expression and release, western blot analysis was used to measure Gag from the producer cell lysate and Gag from the other half of the final washed, resuspended virus used for the infectivity assays (Fig. S9A). As LEN concentration was increased, we observed a correlated decrease in total Gag (used as a proxy for viral protein expression) (Fig. 3C). Despite a 65% reduction in expression at 50 nM LEN compared to DMSO, the corresponding released virus showed only a ∼20% decrease in release at 50 nM LEN compared to DMSO (Fig. 3D). Therefore, the reduction in infectivity cannot be explained completely by the reduction in expression or release.

We hypothesize that the gross morphological effects of LEN on immature assembly and maturation described above are the primary source of the decrease in infectivity observed here. To test this, we performed qPCR analysis of RT products (42). If the virus from cells pretreated with LEN have defective Capsid cores, we expect an early loss of RT products. To ensure that the measured cDNA transcripts corresponded directly with the infectivity measured via flow cytometry, DNA was extracted in parallel from the other half of the same infected cells (Fig. S9A). With increasing concentrations of LEN in the producer cells, we observed corresponding decreases in transcripts for early, intermediate, late, and 2LTR circles in the infected target cells, compared to cells infected with control virus from DMSO treated producer cells (Fig. 3E). In the converse control, target cells treated with 5 nM LEN for 24 h were then infected with effector-naïve virus, which resulted in a reduction of late and 2LTR circle transcripts as has been previously described (11, 12) (Fig. 3E). This data is consistent with our hypothesis that the block to infection caused by LEN pretreatment of producer cells is due to LEN’s disruption of proper lattice formation, given that intact cores are required for efficient reverse transcription (14, 43).

Consistent with our results, others have reported that LEN leads to a decrease of HIV-1 Gag in producer cells (11, 12); however, the source of the reduction is unclear. To address this, we utilized confocal microscopy and transfection of a Gag-mNeonGreen (mNG) fusion protein to visualize localization in cells treated with DMSO or LEN (Fig. 3F and Fig. S10A). In cells treated with DMSO before transfection, Gag-mNG was diffuse throughout the cytoplasm with small puncta localized primarily to the PM (Fig. 3F and Fig. S10B), as reported by many groups previously (44-48). However, increasing concentrations of LEN resulted in increasing amounts of abnormal (mis-localized and larger) Gag-mNG puncta (Fig. 3F and Fig. S10B).

Interestingly, LEN did not appear to affect the total Gag-mNG signal observed by confocal microscopy, which was inconsistent with the western blot measurements in Fig. 3C and previous reports (11, 12). To assess the possibility that the observed larger puncta represented an insoluble Gag-mNG fraction formed by the presence of LEN, we modified our cell lysate extraction protocol to include both non-denaturing conditions (RIPA buffer) or denaturing conditions (8 M urea lysis buffer without SDS) (49). Lysates extracted with RIPA buffer again displayed a corresponding loss of Gag signal from cells pretreated with 50 nM LEN. However, lysates extracted with 8 M urea displayed the same levels of Gag signal across DMSO and LEN treatments (Fig. 3G). This recovery of Gag signal in denaturing conditions is consistent with localization into insoluble aggresomes, which are not dissolved in RIPA buffer, as opposed to LEN decreasing Gag levels in the cell.

We conclude the following: 1) LEN does not reduce Gag expression, 2) LEN does reduce Gag solubility, 3) LEN has a modest effect on virus release (assay dependent), 4) LEN causes assembly defects (premature lattice), and 5) LEN prevents proper maturation. Taken together, these observations demonstrate that LEN blocks infectious virus production.

## Discussion

With the advent of HIV-1 Capsid binding effectors, a better understanding of the dynamics that dictate Capsid conformation, gross morphology, and ligand binding is needed. This is especially true in the context of the dual functionality of CA as both a domain of Gag and mature protein. Here, we demonstrated that LEN affects assembly and maturation by binding the nascent immature lattice, leading to incorporation of the effector into the assembling virus particle and, consequently, to inhibition of multiple downstream steps in the lifecycle.

The effect of LEN on *in vitro* assembled and *in situ* released VLPs was drastically different. In the absence of LEN, VLPs from both systems exhibit an immature lattice. In the presence of LEN, the effect on *in vitro* VLPs was modest, only apparent upon cryo-EM structure determination (Fig. 1B), while *in situ* VLPs were clearly morphologically aberrant (Fig. 2A). *In vitro*, LEN binds to the immature lattice CA_NTD_ similarly to how it binds to the mature lattice FG pocket. LEN occupancy results in an ∼14° rotation around the CA_NTD_ threefold formed by αH2. However, the CA_CTD_ and 6-helix bundle remained unaltered (Fig. 1). These findings are consistent with a previously published structure of the immature lattice of VLPs released from cells, permeabilized, and subsequently treated with LEN (29).

Our *in situ* data suggest that LEN binds to Gag prior to assembly, causing the CA domain to adopt a mature conformation which promotes the assembly of a mature-like CA lattice, i.e. a premature VLP. If this is true, we expect that LEN binding bypasses IP6-dependent immature assembly (23, 31, 32). Consistent with this, IP6 density is weak to absent from the premature and mature lattices produced in the presence of LEN (Fig. 2A). Also, LEN attenuates the impact of IP6 removal from cells via an IPPK-KO (23) (Fig. 2D). However, using a pre-established protocol with “high” concentrations of IP6 (23), our *in vitro* assembly results are not, at first look, in full agreement with this model. However, at lower concentrations of IP6, LEN promoted the formation of mature-like lattice (Fig. 2C). Studies show that total cellular IP6 ranges from 10-40 µM (30), but the available IP6 concentration is likely much lower at 1 to 2 µM (50). Our assembly results are reasonable in the context of this lower and, likely, limiting concentration of available IP6.

Our results are consistent with the observations that LEN stabilizes the hexamer-containing tubular or flatter regions of the lattice and destabilizes the pentamer-containing high curvature regions of the lattice (13, 16, 17). For example, VLPs collected from LEN treated cells all had flat lattice that cryo-EM revealed to contain only CA hexamers (Fig. 2A). Interestingly, we did not observe any CA pentamers in these samples. The CA hexamers had LEN bound but, surprisingly, did not have density corresponding to IP6. The lack of IP6 may also be the reason for the lack of pentamers as we and others have shown that pentamer formation is highly dependent on it (24-26). In addition, the presence of LEN may have prevented pentamers from forming or disrupted them after they formed. In the absence of the pentamer, the Capsid core cannot close and, therefore, is unstable and not infectious. In summary, there may be multiple ways to explain these observations. LEN binding to hexamers flattens and rigidifies the lattice which is unfavorable for pentamers. However, this model does not exclude the possibility that LEN may also bind to pentamers and, by doing so, destroy them. While our results are consistent with those published previously, more work must be done to resolve the underlying mechanisms of LEN induced Capsid disruption.

WT immature assembly results in VLPs that are ∼100-120 nm in diameter. In the presence of LEN, we observed premature (PR-D25A) Gag and mature (PR-WT) CA lattices that result in aberrant VLPs that ranged in size of ∼100-500 nm in diameter (Fig. 2A and Fig. S8A). Native immature budding requires cell-mediated Endosomal Sorting Complex Required for Transport (ESCRT) budding that is dependent on normal virion curvature (51). We predict VLP release in the presence of LEN is independent of ESCRT. However, efforts to demonstrate this via late domain mutations in Gag (52, 53) were inconclusive in the cell-culture system used and should be followed up in future work using alternative systems.

As seen in *in vitro* studies (11, 12, 54) and clinical trials (55, 56), many LEN-resistance mutations have already been identified (57). While only a small percentage of relevant, naturally occurring polymorphisms are in circulation (58, 59), their presence and selection in patients undergoing treatment with LEN necessitate design of second-generation Capsid effectors. The first step in such design is identifying how resistance mutations evade inhibition at the molecular and structural level. Here, we present the putative hydration state of the LEN binding pocket before and after binding as well as the first structure of LEN bound to the RAM M66I (to immature lattice, Fig. 1C). These findings can help in the development of modified moieties to overcome the limitations of LEN in the context of RAMs.

Our results are consistent with what has been reported for LEN, and we propose the following model. In the context of effector-naïve virus infecting effector treated cells, we hypothesize that the limited IP6 pool in the cells renders virus particles susceptible to LEN modulation of the Capsid lattice. We expect that during Capsid trafficking to the nucleus, LEN binds Capsid thereby disrupting the lattice curvature to release vgRNA prematurely or to prevent nuclear entry as has been proposed (15, 16). In the converse situation of effector-treated virus infecting effector-naïve cells, LEN packaged into virus particles during assembly is sufficient for its downstream inhibitory effects (Fig. 4, bottom). The simplest model to explain this notable observation is that LEN predisposes the immature lattice to a flattened premature conformation with limited capacity to bind IP6. A LEN stabilized, flattened lattice would likely lack the flexibility required for canonical ESCRT-mediated scission (51) and would limit ESCRT induce curvature (60). Thus, virus release, in part, would require that patches of lattice be sloughed off the producer cells. The virus particles that do form, however, are enlarged and contain reduced IP6. Following proteolytic processing, Capsid cores would be unable to close due to a lack of IP6 as well as to LEN binding to the FG pocket, both of which prevent proper pentamer formation. Upon viral fusion with a target cell, we expect that the little core closure that does occur is antagonized by interactions with host proteins such as Cyp-A (15), TRIM-5α (61, 62), FEZ1 (63, 64), and many others (65), thereby exerting pressures on the core. This pressure results in premature release of the Capsid contents, rendering RT incapable of reverse transcribing the vgRNA and, thus, halting infection. Of the few intact Capsids that are able to traffic to the nucleus, the already bound LEN blocks interactions with the Nups and CPSF6 (11, 12, 66) providing a final block to infection.

**Figure 4.**
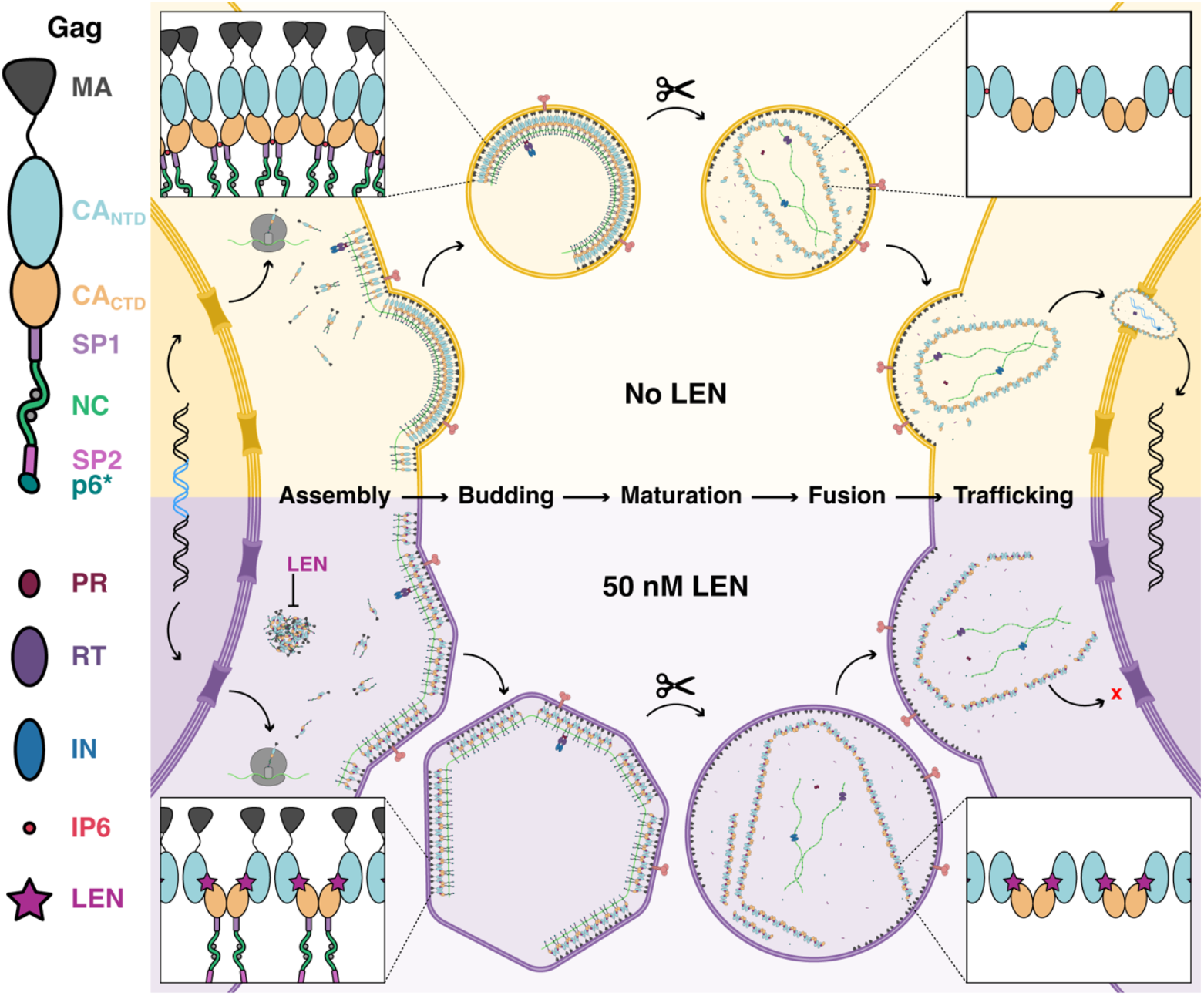
Summary of results and hypotheses. Key: Gag, Pol proteins, IP6, and LEN. Main, top: schematic of the retroviral life cycle without LEN. Main, bottom: schematic of the retroviral life cycle in the presence of 50 nM LEN (pre-treated producer cells). In the presence of LEN, we observed putative aggresomes in the cytoplasm. Virus that does assemble at the plasma membrane, presumably, lacks curvature as inferred from the flat facets observed in the released VLPs. These VLPs also exhibited substantially diminished IP6, which contributes to the lack of pentamers, and is consistent with the lack of curvature and failure to close following maturation. Upon fusion, the aberrant mature cores are unable to sustain proper reverse transcription of the vgRNA and are impaired in their ability to dock at the nuclear pore.

## Materials and Methods

### Protein purification and in vitro assembly

*E. coli* vectors for *in vitro* HIV-1 CA expression were previously described (26). *In vitro* protein expression purification, and assembly were previously described (23, 26). Briefly, HIV-1 CASPNC and CA proteins were expressed and purified from NiCo21(DE3) competent *E. coli*. Purified proteins were concentrated to ∼2 mg/mL and ∼20 mg/mL for CASPNC and CA respectively, flash frozen, and stored at -80°C.

*In vitro* assembly was performed as previously described for immature and mature VLPs and CLPs (23) but with the following alterations corresponding to LEN addition. Following CLP assembly, LEN in DMSO was added to the reaction at the indicated concentration (see Table S1). For CASPNC assembly LEN addition was added before, during, or after assembly as described below. Addition before: protein was pre-incubated with LEN for ∼2 hours prior to the addition of IP6 and nucleic acid and initiation of assembly via dialysis (23). Addition during: protein, IP6, nucleic acid, and LEN were mixed prior to initiating assembly via dialysis. Addition after: two or more hours after assembly, LEN was added to VLPs. For *in vitro* CLP assembly, assembly buffer was warmed to 37 °C followed by addition of CA and incubated for 15 min. All assembly reactions were prepared for negative stain transmission electron microscopy (23) and imaged on one of the following microscopes (FEI Tecnai 12 BioTwin TEM, FEI Morgagni TEM, ThermoFisher Talos L120C TEM, or DeLong LVEM25) for sample abundance and morphology.

### Cryo-EM imaging and data processing

Screening, sample preparation, and data collection are described here (25, 26) as well as in Table S1 and Figs S2, S5, and S7. *In vitro* assembled and *in situ* purified VLPs were vitrified using standard methods (19, 67). Micrographs were collected at 63,000x or 105,000x in super resolution mode on a 200 kV Talos Arctica TEM or on a 300 kV Titan Krios TEM (ThermoFisher) respectively with SerialEM (68). Both microscopes were equipped with a K3 direct detector (Gatan) and a BioQuantum energy filter (Gatan).

SPA image processing was performed as previously described (25, 26) using the pipeline indicated in Table S1 and Figs. S2, S5, and S7. Briefly, SPA image processing was done in RELION v4.0.1-5.0 (69) and CryoSPARC v4.7.0 (70). Motion correction and CTF estimation were carried out using MOTIONCOR2 (71) and GCTF (72). Maps and models were visualized with IMOD (73), Chimera (74), and ChimeraX (75).

### Model refinement

The preliminary models for the LEN-bound CA (PDB: 8G6M) (26) and Gag (PDB: 5L93) (20) assemblies were refined using molecular dynamics flexible fitting (MDFF) (76, 77). Briefly, the preliminary models and parameterized LEN were prepared for MDFF by assigning protonation states at pH 7.0, solvating the system, and adding Na^+^ and Cl^-^ ions around the protein as well as the bulk solvent to 150 mM NaCl via VMD (78). The MDFF ready system was then refined into the cryo-EM density using resolution exchange MDFF (ReMDFF) (76). Agreement of the protein sidechains was evaluated via local cross-correlation, and the coordinates of the best fitting sidechains were used to create a ReMDFF-derived model. See *SI Methods* for more details. All MD simulations were performed in the NAMD v3.0.1 (79) simulation engine using CHARMM36m (80) force-field parameters for proteins, water molecules and ions, CGenFF (81) derived parameters for IP6, and QM-optimized CHARMM parameters for LEN, as detailed in the *SI Methods*.

### Force-field parameterization of LEN

To perform molecular dynamics flexible fitting and minimizations of the LEN-bound CA structures, we derived CHARMM force field parameters for LEN using the Force Field Toolkit (ffTK) (82) plugin in VMD (78) and the CHARMM general force field (CGenFF) (81, 83) parameterization approach. First, we performed DFT geometry optimization of the Lenacapavir coordinates from the co-crystal structure of LEN bound to an HIV-1 capsid hexamer (PDB ID: 6VKV) (12). Once optimized, we derived an initial set of force field parameters for LEN by analogy to CGenFF (83) parameters for small molecules. From the initial set of parameters, parameters with high penalties (>10) were identified (Fig. S3A) and targeted for further optimization. To reduce the computational cost of these calculations, we utilized a “divide and conquer” approach as recommended by the MacKerell lab (83) for parameterizing large compounds, dividing Lenacapavir into smaller molecule fragments containing the regions of interest for parameter optimization and capping terminal bonds with hydrogen (Fig. S3B). To parameterize partial atomic charges of water-accessible atoms, we placed a water molecule 2 Å from the atom of interest forming a hydrogen bond and we optimized the distance and orientation of the water molecule at QM level with all other degrees of freedom fixed. Bond and angle parameters were optimized from a QM vibrational analysis. Finally, to parameterize dihedral angles, we performed potential energy scans for each dihedral, and we fit the parameters in MM dihedral potential energy functions to match the potential energy surface measured from the QM scans (Fig. S3C). Once the optimized parameter set was obtained for the molecule fragments, the fragment parameters are merged into the full molecule. See *SI Methods* for more details.

### VLP release, purification, and western blot analysis

Protease inactivating mutation D25A (84) and Gag-M198I (CA-M66I) (11, 12) were made in the proviral vector HIV-1^Δenv^ (40) using In-Fusion cloning. All vectors were verified by sanger sequencing. 293FT cells were seeded in 15 cm cell culture dishes in complete media with either DMSO or LEN at the indicated concentrations for cryoEM (Fig. S5 and S7). 24 h post-seed, cells were transfected with proviral vector using Lipofectamine 3000. VLPs were purified as previously described (19) with some details here. 48 h post-transfection media was centrifuged to remove cell debris and the resulting supernatant was centrifuged through a 20% sucrose-PBS cushion and the resulting viral pellet was resuspended. Subsequently, the cleared VLPs were subjected to density gradient sedimentation: OptiPrep-PBS (6% to 18% in 1.2% steps and a 35% stop) and 200k x g for 1.5 h at 4°C. Fractions were collected and subjected to western blot analysis. CA-containing fractions were pooled, diluted with final suspension buffer (25 mM Tris pH7.4 at RT and 150 mM NaCl), and pelleted to concentrate the VLPs for cryo-EM. WT and Gag-M198I (CA-M66I) virus release assays from 293FT and 293FT-ΔIPPK cells were performed as previously described (23). For western blots, the antibodies used are as follows: RbαHIV-p24 (NIH HIV Reagent Program/BEI Resources), AlexaFluor680 conjugated GtαRb-IgG, and AlexaFluor790 conjugated MsαGAPDH. Probed membranes were imaged using a Li-COR Odessey or Azure600.

### Transductions, flow cytometry, and qPCR

Workflow of VLP purification for infectivity is shown in Fig. S9A and described briefly here. 293FT cells were seeded (4.5E5 cells per well) with 2 mL of complete media in 6-well format and treated with DMSO (negative control) or DMSO with 0.05, 0.50, 5.00, or 50.0 nM LEN. 24 h post-seed, cells were transfected with HIV^ΔEnv^ and Vesicular Stomatitis Virus glycoprotein (85) vectors using Lipofectamine 3000. 48 h post-transfection, VLP containing media was collected and cell debris removed via centrifugation. To wash the VLPs of residual LEN, supernatant was pelleted through a 20% sucrose-PBS cushion, resuspended in PBS, and pelleted again before final resuspension in PBS for subsequent infectivity assays. The transfected cells were harvested and lysed in RIPA buffer. The cell lysate and a fraction of the washed VLPs were analyzed via western blot to measure CA levels as a proxy for virus production and release (23).

Cells for the infectivity assay were seeded and treated with 5.00 nM LEN (positive control) (11, 12) or DMSO (experimental). 24 h post-seed, LEN treated cells were transduced with washed VLPs from DMSO treated producer cells, and DMSO treated cells were transduced with washed VLPs from LEN treated producer cells. 6 h post-transduction, cells were lifted. Half of the cells were used for DNA extraction to measure early RT transcripts (42), and half were replated. 48 h post-transduction, cells were collected again. Half were used for DNA extraction to measure intermediate, late, and 2LTR circle transcripts (42). The other half were fixed in 5% PFA-PBS, washed, and analyzed for fluorescence using an Attune Flow Cytometer to measure transduction. The number of transduced cells gated by green fluorescence (Fig. S9B) in the LEN treated samples were normalized to the number of transduced cells in the DMSO control and presented as percent infectivity.

### Data analysis

A Plasmid Editor (APE) was used for plasmid cloning and sequence analysis (86). Band densities from western blot images were analyzed using GelAnalyzer (87). Flow cytometry data were analyzed using the Attune Collection and Analysis software (ThermoFisher). qPCR data was analyzed via the Design and Analysis Suite (Applied Biosystems by ThermoFisher). Raw values for all data points were inputted into Excel and normalized data exported to CSV format for statistical analysis and graph generation via R (v4.5.1) (88) in RStudio (v2025.5.0.496) (89). Images for cryo-EM maps and atomic models were generated via ChimeraX (75).

### Data availability

Cryo-EM maps and associated atomic coordinates have been deposited into the World-Wide Protein Data Bank (Electron Microscopy Data Bank and Research Collaboratory for Structural Bioinformatics Protein Data Bank): EMD-75781 and PDB-11KU (*in vitro* CASPNC-WT immature with 0 μM LEN at 300 kV), EMD-75782 and PDB-11KV (*in vitro* CASPNC-WT immature with 50 μM LEN at 300 kV), EMD-75783 (*in vitro* CASPNC-WT immature with 50 μM LEN at 200 kV), EMD-75784 and PDB-11KW (*in vitro* CASPNC-M66I immature with 50 μM LEN at 200 kV), EMD-75785 (*in situ* CA-WT PR-D25A immature with 0 nM LEN at 300 kV), EMD-75786 (*in situ* CA-WT PR-D25A immature with 10 nM LEN at 300 kV), EMD-75787 (*in situ* CA-WT PR-D25A premature with 10 nM LEN at 300 kV), EMD-75788 and PDB-11KX (*in situ* CA-WT PR-D25A premature with 50 nM LEN at 300 kV), EMD-75789 (*in situ* CA-M66I PR-D25A immature with 50 nM LEN at 200 kV), EMD-75790 (*in situ* CA-WT PR-WT mature with 50 nM LEN at 300 kV), and EMD-48430 and PDB-9MNM (*in vitro* CA-WT mature with 500 μM LEN at 300 kV).

QM-derived CHARMM force-field compatible parameters for Lenacapavir are available in the Perilla group’s HIV inhibitor force-field repository: https://github.com/Perilla-lab/HIVff.

## Acknowledgments

This work was supported by the National Institute of Allergy and Infectious Diseases under awards R01AI147890, R56AI189254 and Collaborative Development Program 5U54AI15047209 to R.A.D., and R01AI178846 and U54AI170791 to J.R.P. A portion of this work was also supported by U54AI170855 and R37AI150479. Cryo-EM data collection performed at the Cornell Center for Materials Research supported by the National Science Foundation award DMR-1719875. A portion of this research was supported by NIH grant R24GM154185 and performed at the Pacific Northwest Center for Cryo-EM (PNCC) with assistance from V. Rayaprolu. This work used the Stampede3 supercomputer at the Texas Advanced Computing Center (TACC) through allocation MCB-170096 from the Advanced Cyberinfrastructure Coordination Ecosystem: Services and Support (ACCESS) program, which is supported by National Science Foundation awards #2138259, #2138286, #2138307, #2137603, and #2138296. We acknowledge computational support through the Delaware Advanced Research Workforce and Innovation Network (DARWIN) at the University of Delaware. We also thank Volker M. Vogt and Stefan G. Sarafianos for helpful discussions.

