## Supplemental Information for "Lenacapavir prevents production of infectious HIV-1 by abrogating immature virus assembly"

### Supplemental Methods

*Model refinement:*

The preliminary models for the LEN-bound CA (PDB: 8G6M) (1) and Gag (PDB: 5L93) (2) assemblies were refined using molecular dynamics flexible fitting (MDFF) (3, 4) through the following procedure: First, protonation states at pH 7.0 for protein residues in all preliminary models (CA in complex with LEN, CA in complex with IP6 and LEN, CASP1 in complex with IP6 and CASP1 in complex with IP6 and LEN) were assigned using the PDB2PQR (5) software (v3.6.1). For the models involving LEN-bound mature CA assemblies, the coordinates of the 2-[(2S,4R)-5,5-difluoro-9-(trifluoromethyl)-7,8-diazatricyclo[4.3.0.02,4]nona-1(6),8-dien-7-yl]acetamide moiety of LEN were obtained by aligning the LEN coordinates from the co-crystal structure of LEN bound to an HIV-1 capsid hexamer (PDB ID: 6VKV) (6). Furthermore, to prepare the models for MDFF (3, 4), the protein-ligand systems were solvated in a periodic box with TIP3P (7) water molecules, using the solvate plugin in VMD (8). To simulate physiological conditions, we added $Na^+$ and $Cl^-$ ions around the protein and in bulk solvent to a 150mM NaCl concentration using the cionize and autoionize VMD plugins. In addition, to increase the simulation timestep in MDFF to 4.0 fs, we redistributed the mass of heavy atoms and bound hydrogens in protein and ligands used the hydrogen mass repartition (HMR) (9) scheme. MDFF-ready systems consisted of ~600,000 atoms for mature CA and ~574,000 atoms for immature CASP1 assemblies, including solvent and ion atoms.

The coordinates of the protein and ligand atoms in MDFF-ready systems were then refined into the cryoEM density utilizing resolution exchange MDFF (ReMDFF) (3). In MDFF, a molecular dynamics simulation is performed where the position of the protein and ligand heavy atoms are coupled to a grid-based potential, biasing the movement of these atoms to fit in a reference density map. In ReMDFF, multiple replicas of MDFF are run in parallel using maps of sequentially higher resolutions to allow for fitting of the global characteristics of the molecular structure as well as the atomistic details. ReMDFF has been shown to converge faster for high-resolution cryoEM densities in comparison to direct MDFF (3), since it allows higher conformational diversity, and prevents the system from becoming trapped in a local minima when fitting high-resolution density maps.

To create the biasing maps for ReMDFF, we used the experimental cryoEM density maps (Fig. S2) and generated 7 lower resolution maps by applying sequential Gaussian smoothing filters (σ=0.5 Å) with the voltool command in VMD (8). In total, we performed 8 ReMDFF replicas with progressively higher resolution densities, up to the experimental resolution, sampling 20ns per replica with a coupling grid scaling factor of 0.3 a.u. Replica exchanges between neighboring replicas were attempted every 0.2 ps and were accepted based on the Metropolis criterion, as implemented in Hamiltonian replica exchange (10).

Following ReMDFF, the agreement of the protein sidechains was evaluated via local cross-correlation, and the coordinates of the best fitting sidechains for each resolution were utilized to create a ReMDFF-derived model. The ReMDFF process may distort the length of molecule bonds due to the additional density-based grid potential applied to the protein and ligand atoms. Therefore, to reduce this strain while maintaining proper sidechain fitting to the density, we performed successive energy minimizations with decreasing coupling grid scaling factors (from 1.0 a.u. to 0 a.u. in 0.1 a.u. steps) until the density-guided potential was turned off. The minimized structures agree with the cryoEM density and conserve the proper bond-lengths in molecular mechanics force-fields.

All ReMDFF calculations were performed on an ensemble with constant number of atoms, pressure and temperature (NPT). Pressure was maintained at 1 atm via the Nose-Hoover barostat with period and decay parameters of 200 ps and 100 ps, respectively, and temperature was maintained at 310 K using a Langevin thermostat with damping constant of 5 $ps^{-1}$. A timestep of 4.0 fs was used, enabled by the HBR scheme during system preparation. Nonbonded interactions were calculated with a 12 Å cutoff and 10 Å switching distance for short-range interactions and long-range electrostatic interactions were computed every 8.0 fs using the particle mesh Ewald algorithm (11) with a 1 Å grid spacing. All MD simulations were performed in the NAMD v3.0.1 (12) simulation engine using CHARMM36m (13) force-field parameters for proteins, water molecules and ions, CGenFF (14) derived parameters for IP6, and QM-optimized CHARMM parameters for LEN, as detailed in the following section.

*Force-field parameterization of Lenacapavir:*

Molecular mechanics (MM) potential energy functions require empirical parameters for every bond, angle, and dihedral in the molecule as well as Van der Walls parameters and charges for every atom. To perform molecular dynamics flexible fitting and minimizations of the LEN-bound CA structures, we derived CHARMM force field parameters for LEN using the Force Field Toolkit (ffTK) (15) plugin in VMD (8) and the CHARMM general force field (CGenFF) (14, 16) parameterization approach.

First, we performed DFT geometry optimization of the Lenacapavir coordinates. After the drug geometry was optimized, we derived an initial set of force field parameters for LEN by analogy to CGenFF (16) parameters for small molecules. From the initial set of parameters, parameters with high penalties (>10) were identified (Fig. S3A) and targeted for further optimization. To reduce the computational cost of these calculations, we utilized a “divide and conquer” approach as recommended by the MacKerell lab (16) for parameterizing large compounds, dividing Lenacapavir into smaller molecule fragments containing the regions of interest for parameter optimization and capping terminal bonds with hydrogen (Fig. S3B).

The CHARMM forcefield parameterization philosophy emphasizes reproducing QM interactions with water molecules. Thus, to parameterize partial atomic charges of water-accessible atoms, we placed a water molecule 2 Å from the atom of interest forming a hydrogen bond and we optimized the distance and orientation of the water molecule at QM level with all other degrees of freedom fixed. The atom-water molecule distances, angles, interaction energies, and dipole moment are then used in the ffTK (15) to fit and optimize partial atomic charges. Bond and angle parameters were then parameterized from QM vibrational spectra of the molecule, by comparing and the QM and MM potential energy surfaces calculated from the respective Hessian matrices.

Finally, for parameterizing dihedral angles, we performed dihedral energy scans for each dihedral, and we fit the parameters in MM dihedral potential energy functions to match the potential energy surface measured from the QM scans (Fig. S3C). For the dihedral angle associated with the peptide bond in Lenacapavir, we assigned dihedral parameters by analogy to the observed parameters for Valine in the CHARMM36m (13) force-field for proteins. The total root mean square difference between cumulative QM and MM energy profiles for all dihedrals after our parameterization is 4.4 kcal/mol.

Once the optimized parameter set was obtained for the molecule fragments, the fragment parameters are merged into the full molecule. The charge of the added hydrogen atoms was transferred to their bonded heavy atom, ensuring charge conservation of the full molecule. All quantum mechanics calculations were performed using Gaussian 16 (17) using the B3LYP functional with the 6-31G* basis set and employing implicit water solvent reaction field via the integral equation formalist of the polarizable continuum model (iefpcm).

**Supplemental Figures, Tables, and Legends**

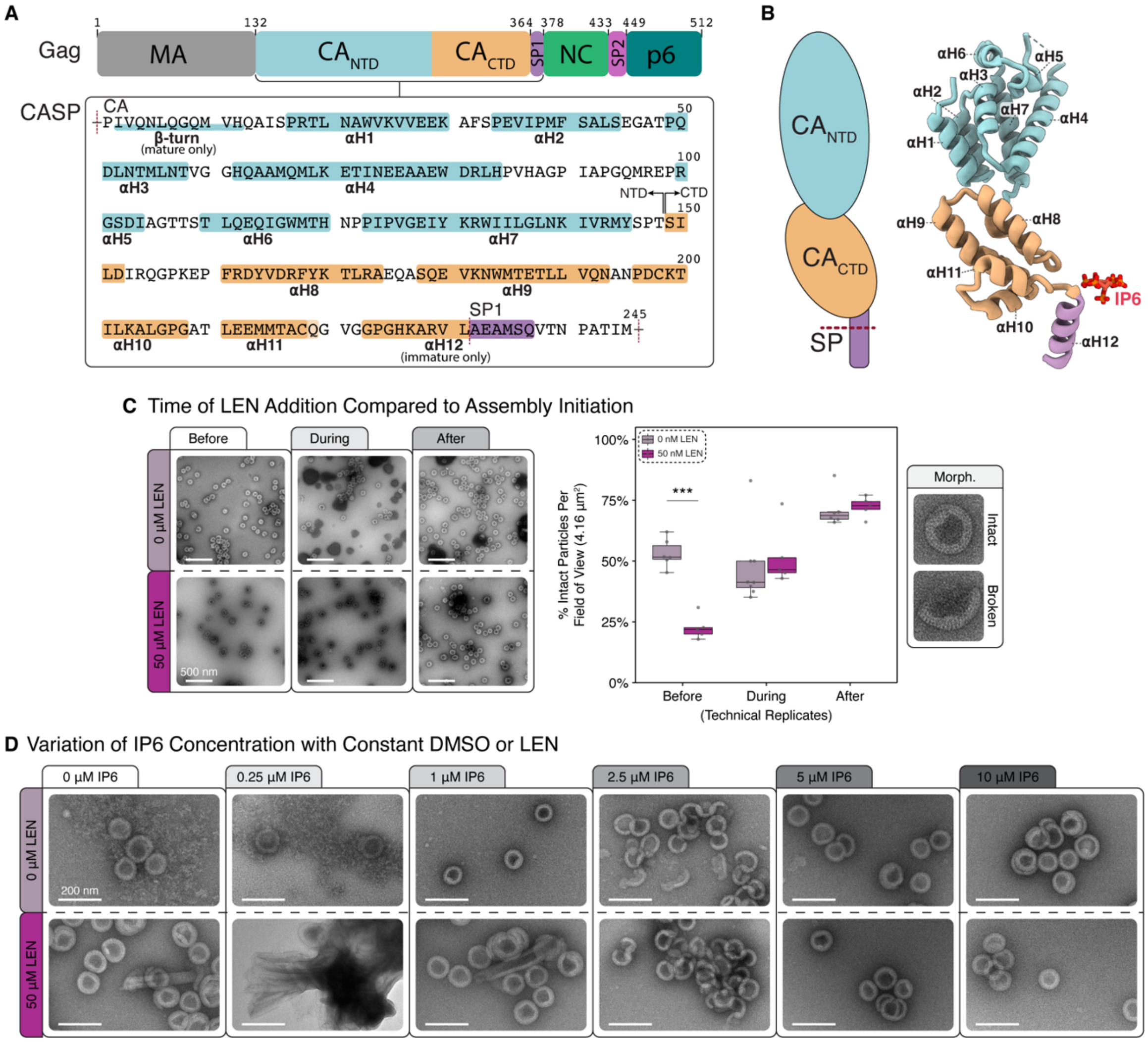


**Figure S1.** *Protein key and biochemical analysis of in vitro assembled immature lattice.* **A)** Top: Schematic of the HIV Gag structural protein and corresponding domains. Gag is numbered from the starting Met of the MA domain. Bottom: Amino acid sequence of CASP. For the purposes of this paper and consistency with current published structures, CASP is numbered from the starting Pro of the CA domain. β-turn (present only in mature lattice/half highlighted) and α-helices are denoted by colored highlighting. The six-helix bundle (αH12) is formed by amino acids from both the CTD and SP. For simplification of maps and diagrams αH12 will be singly colored as lavender. **B)** Left: Cartoon highlighting the domains of the CASP monomer of immature lattice. Right: The tertiary structure of the CASP monomer depicted in ribbon form. **C)** Left: Example nsTEM images of assembled CASPNC with the addition of DMSO or 50 µM LEN 30 min before (left), during (middle), or after (right) initiation of assembly by dialysis with 10 µM IP6 and GT25 oligo. Right: Quantification of intact or broken particles (Technical replicates, n = 6, Student's T-Test, *** P<0.001). **D)** Example nsTEM images of assembled CASPNC with GT25 oligo, the addition of DMSO or LEN during initiation of assembly, and increasing IP6 concentrations.

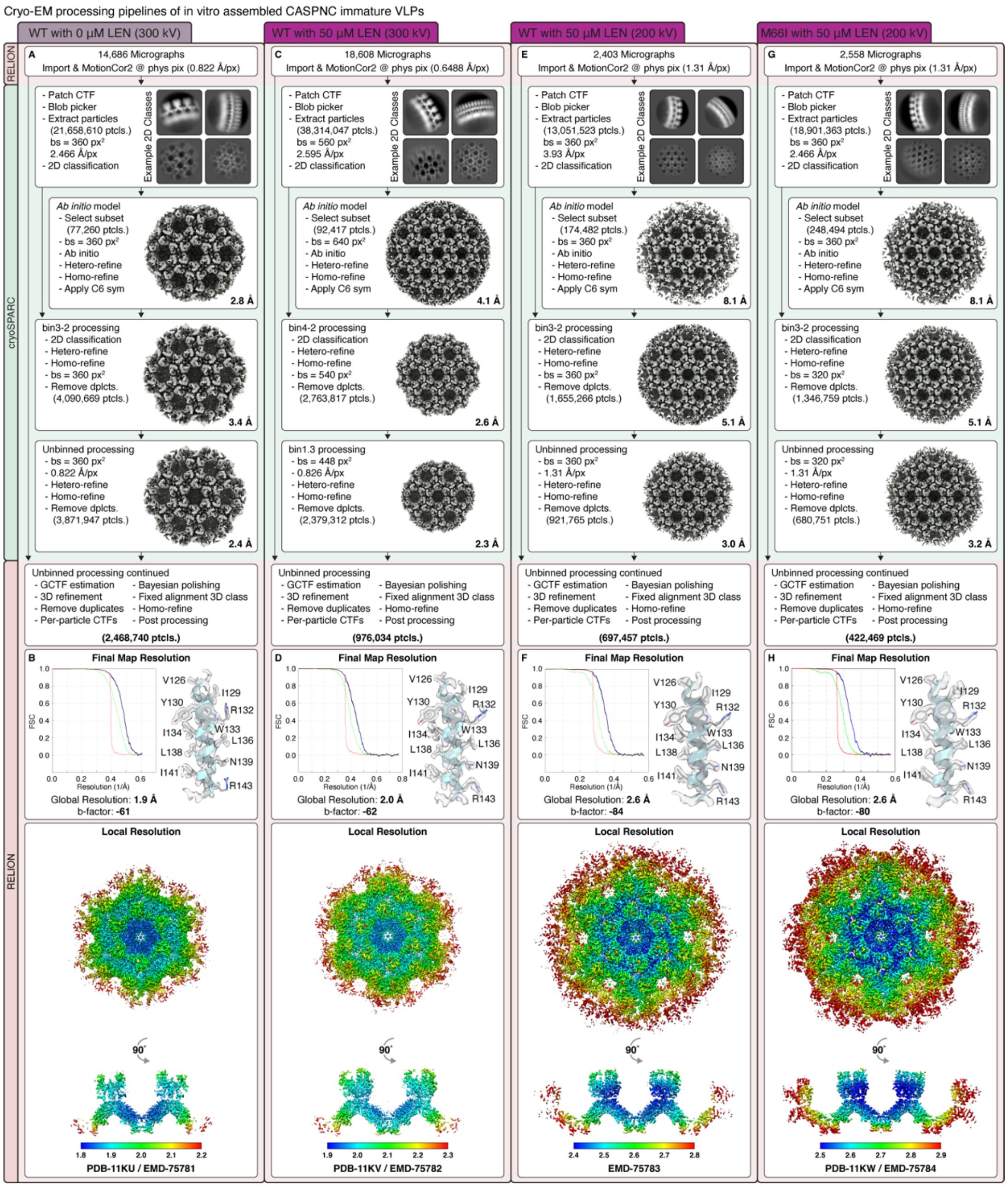


**Figure S2.** *Processing pipeline for in vitro assembled immature protein lattice.* **A)** General processing pipeline for immature lattice from WT CASPNC assemblies with 0 μM LEN. **B)** FSC curve, example map and model of αH7 (CA-numbering), local resolution map of the final refinement (top and side views), and PDB and EMDB accession numbers. **C-D)** As in **A-B** but for assemblies with 50 μM LEN (1:1 concentration of CASPNC:LEN). **E-F)** As in **C-D** but for assemblies collected at 200kV for comparison. **G-H)** As in **E-F** but for CASPNC M66I assemblies with 50 μM LEN collected at 200kV.

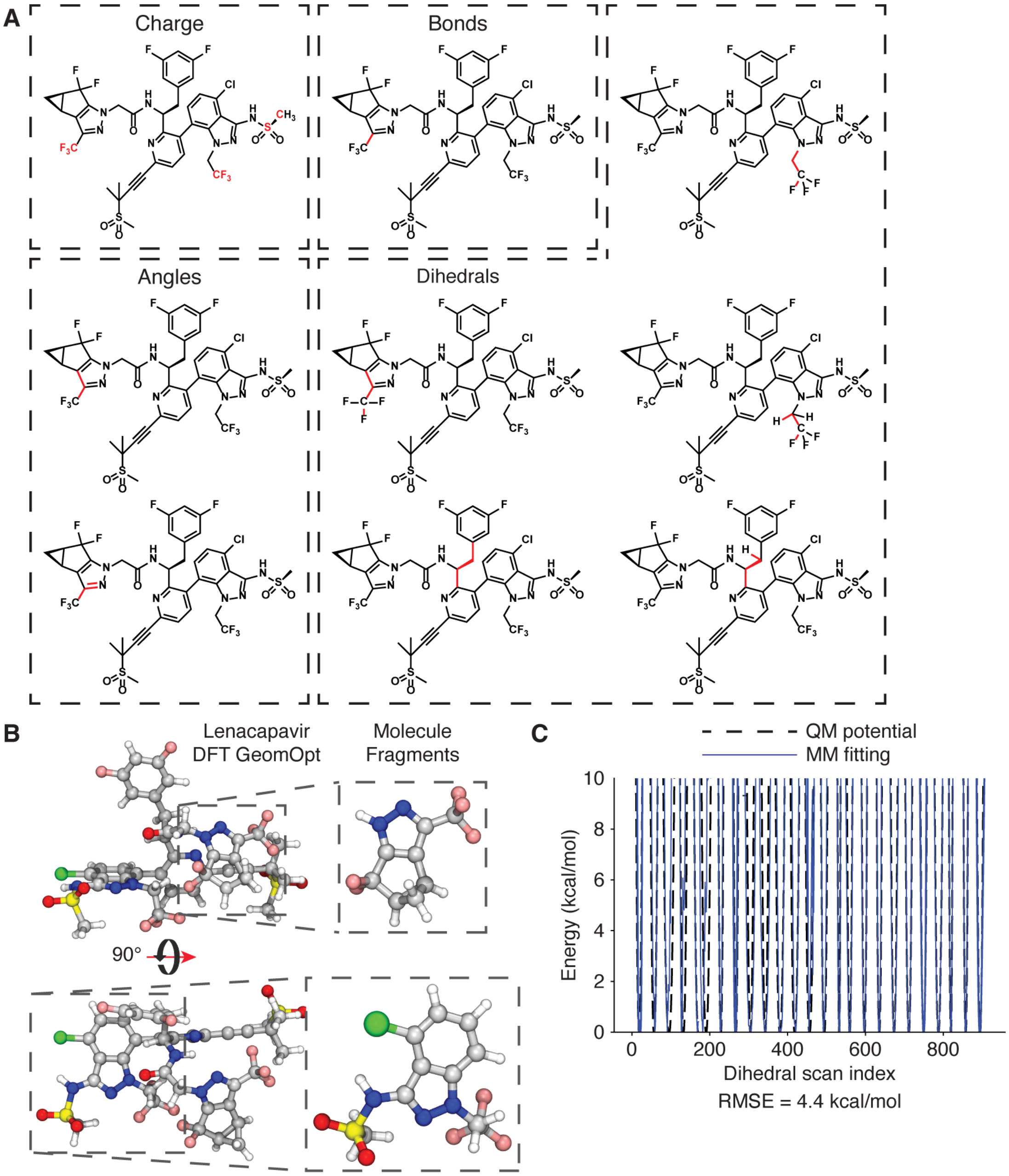


**Figure S3.** *Force-field parameterization of Lenacapavir.* **A)** Chemical structure of LEN overlayed with select high-penalty CHARMM parameters that were optimized via QM calculations (in red). Parameters are divided by type as charge, bond, angle and dihedral parameters. **B)** QM optimized geometry for LEN (left) and the two molecule fragments (right) that were utilized to optimize the high penalty parameters. **C)** Parameterized molecular mechanics potential (blue) and the corresponding quantum mechanics calculated potential for LEN conformations along multiple dihedral scans at B3LYP/6-31G* level of theory (black). The rms difference between both energy profiles is 4.4 kcal/mol.

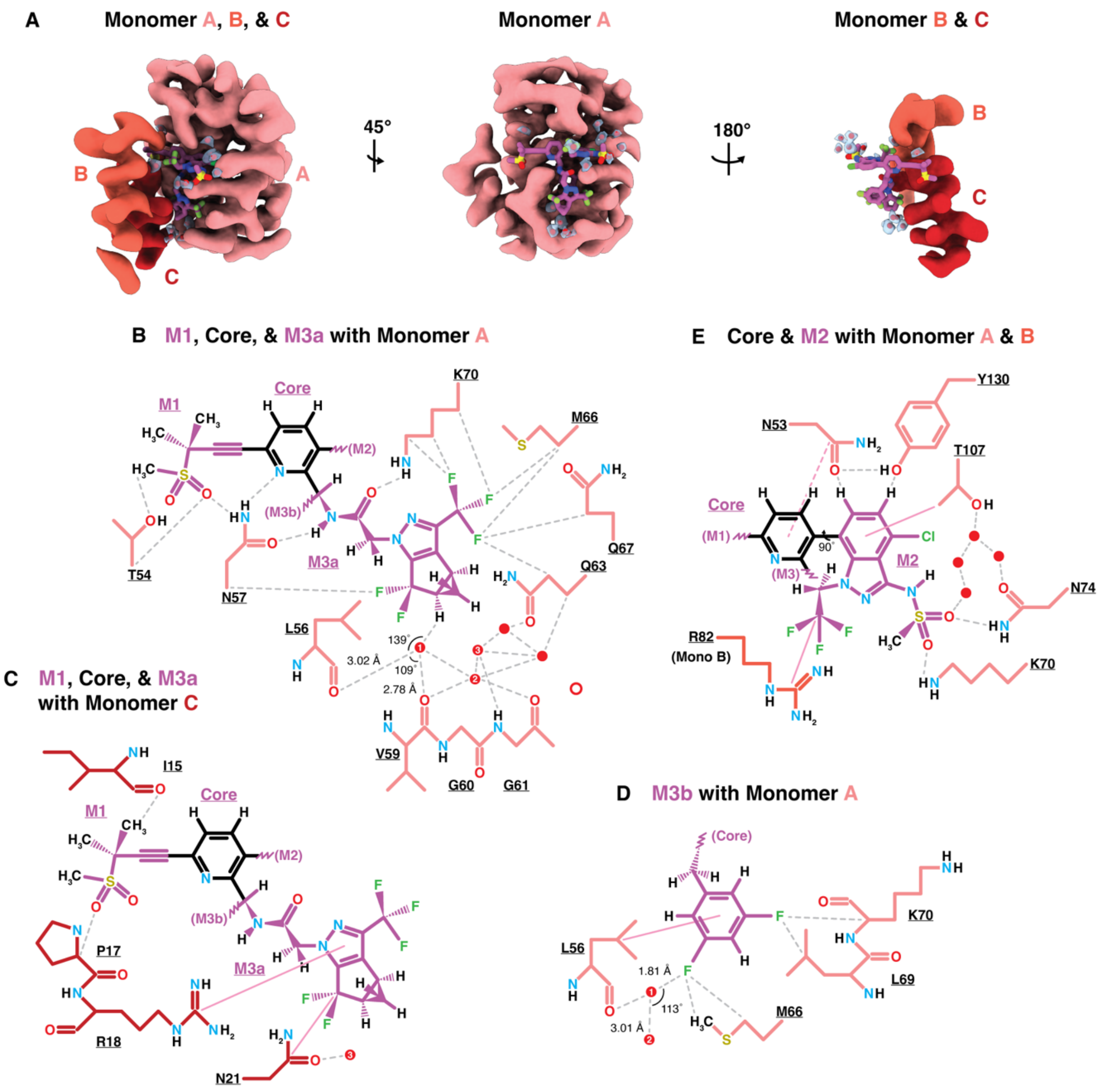


**Figure S4.** *Additional analysis of LEN binding to in vitro assembled immature (CASPNC) protein lattice.* **A)** Map and model depicting the quaternary structure of the LEN binding pocket with modeled waters (α-helices have been lowpass filtered to 4 Å, LEN molecule is shown in purple, and cryo-EM waters determined from the 1.9 Å map are in red/blue). **B-E)** Select 2D interaction models displaying hydrogen bonding (dashed grey) and pi-stacking (solid pink) interactions of amino acids with LEN and waters (red circles). Waters are numbered if they appear in more than one panel. Bond distances and angles for water #1 are indicated.

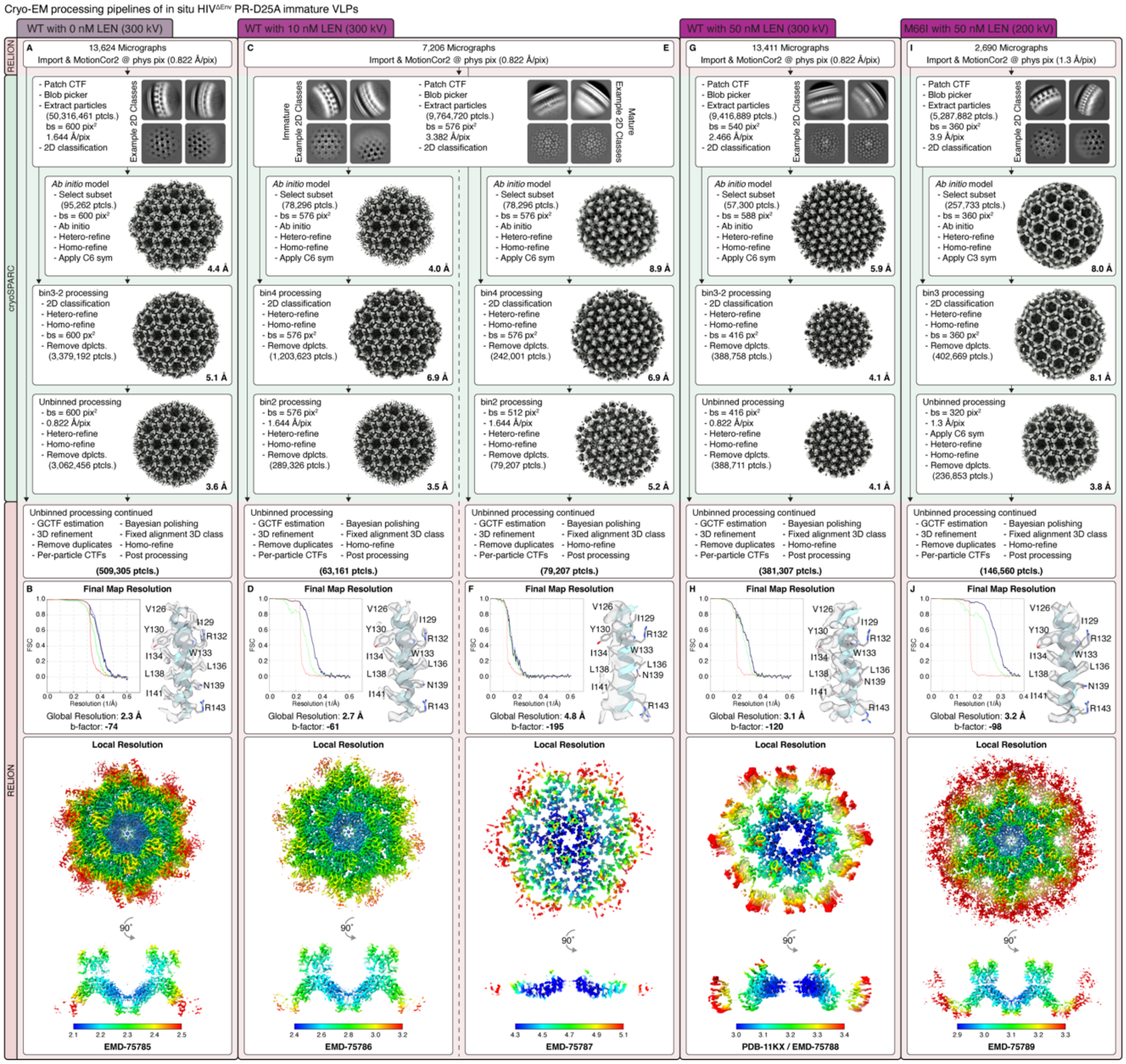


**Figure S5.** *Processing pipeline for in situ VLPs released from cells without or with LEN treatment.* **A)** General processing pipeline for WT-CA PR-D25A VLPs from cells pretreated with 0 nM LEN. **B)** FSC curve, example map and model of αH7 (CA-numbering), local resolution map of the final refinement (top and side views, immature lattice), and PDB and EMDB accession numbers. **C-D)** As in **A-B** but with 10 nM LEN (immature lattice). **E-F)** As in **A-B** but with 10 nM LEN (mature lattice). **G-H)** As in **A-B** but with 50 nM LEN (mature lattice). **I-J)** As in **A-B** but for M66I-CA PR-D25A VLPs from cells pretreated with 50 nM LEN (immature lattice).

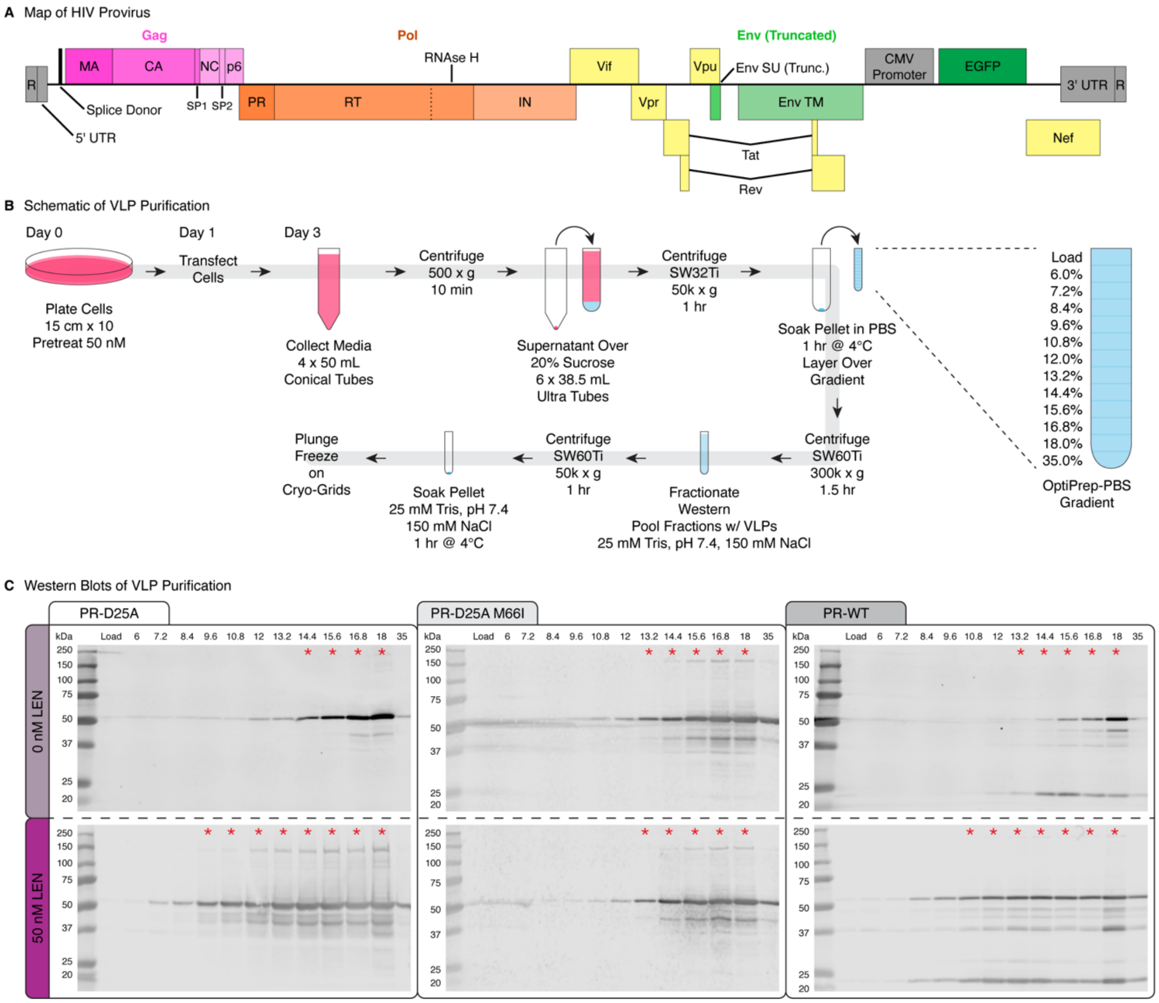


**Figure S6.** *In situ VLP purification.* **A)** Gene map of HIV-1 provirus used to make VLPs. **B)** Schematic of VLP purification from 293FT cells. **C)** Western blots of OptiPrep purification gradients for PR-D25A, M66I PR-D25A, and PR-WT produced in the presence of DMSO or 50 nM LEN. OptiPrep concentrations are shown above the corresponding fractions. Red asterisks indicate sample fractions collected for final purification.

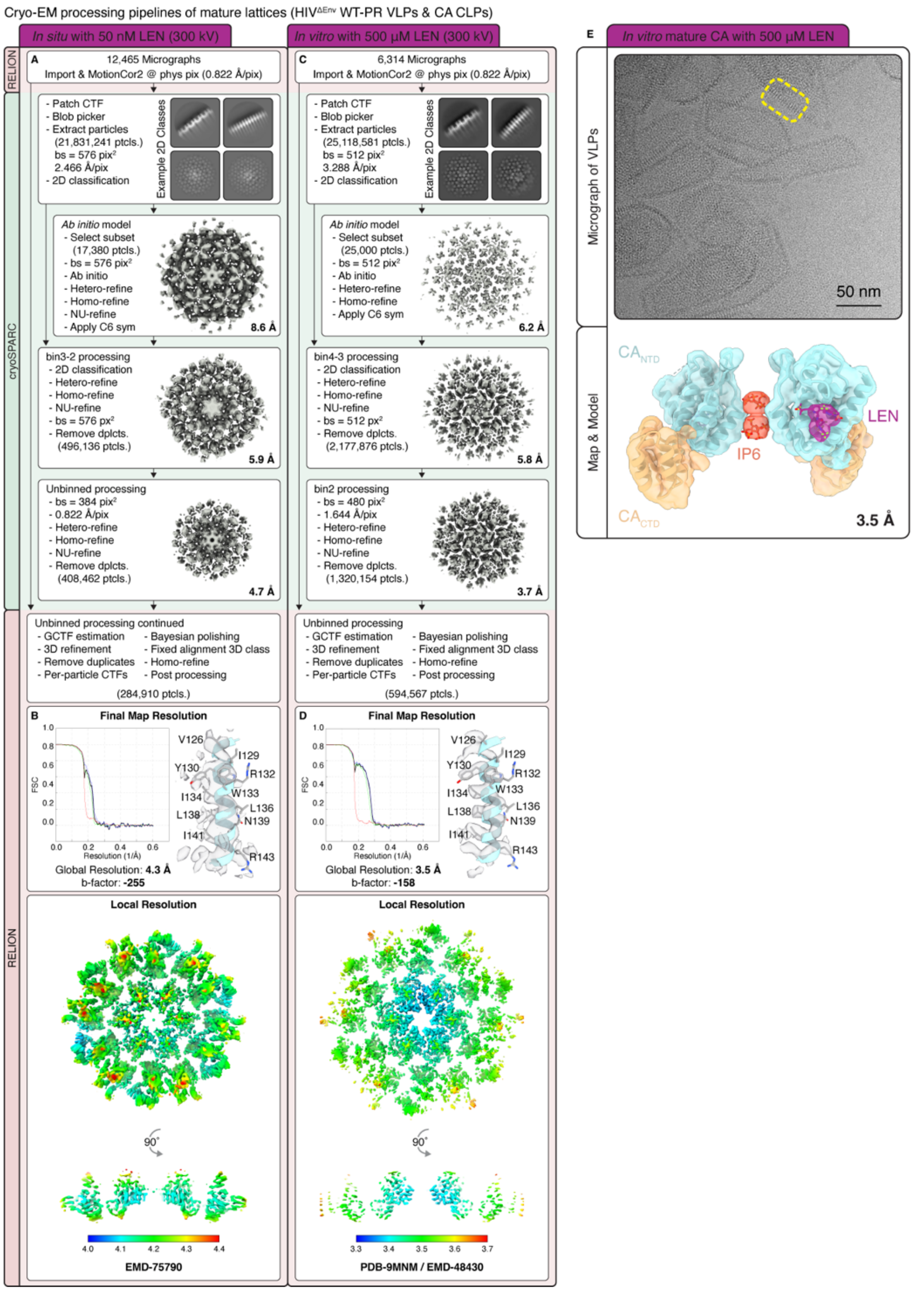


**Figure S7.** *Processing pipeline for mature lattice from VLPs and CLPs.* **A)** General processing pipeline for PR-WT *in situ* VLPs from cells pretreated with 50 nM LEN. **B)** FSC curve, example alpha helix with model, local resolution map of the final refinement (top and side views, mature lattice), and PDB and EMDB accession numbers. **C-D)** As in **A-B** but for *in vitro* CLP assemblies with 500 µM LEN (1:1 concentration of CA:LEN). **E)** Top: Example micrograph of *in vitro* CLPs assembled and then treated with 500 µM LEN. Bottom: Side view of the atomic model within the determined cryo-EM map.

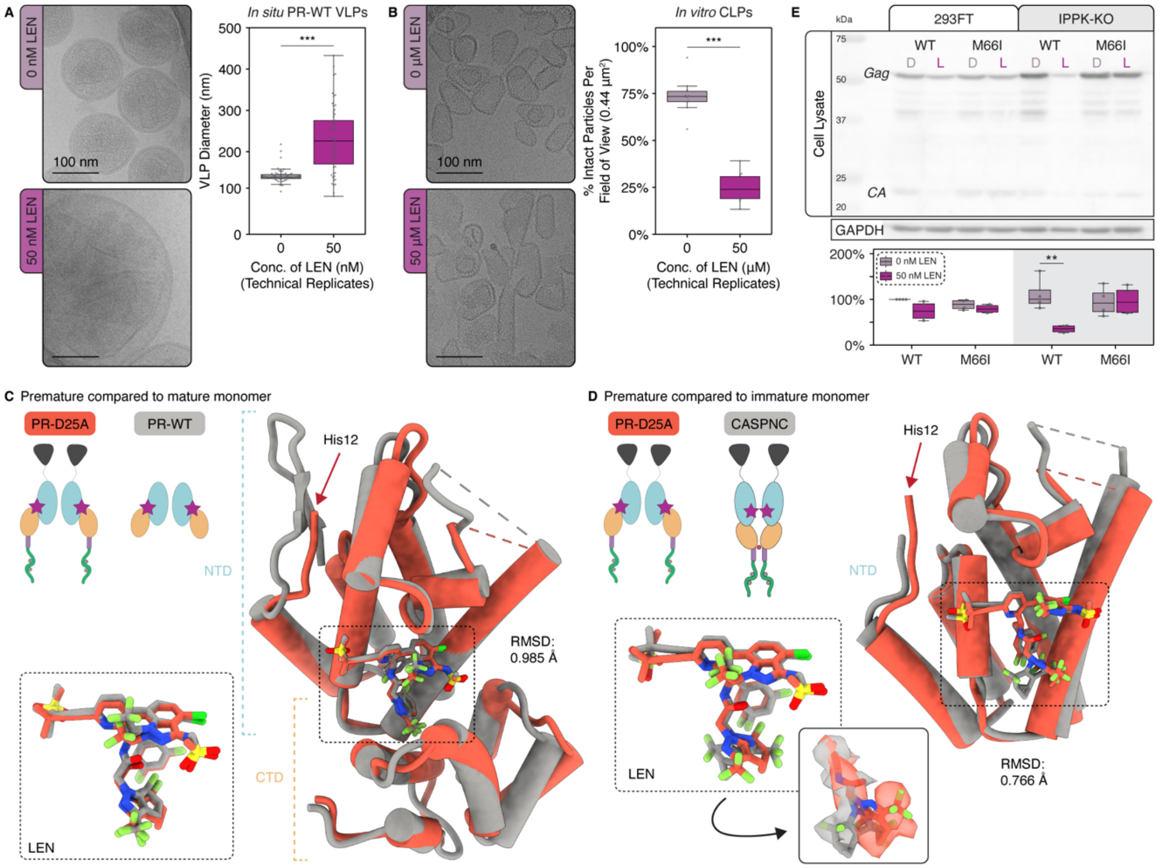


**Figure S8.** *Comparison and analyses between premature and mature lattices.* **A)** Representative micrographs (left) of PR-WT *in situ* VLPs from cells pretreated for 24 hrs with either 0 nM (top) or 50 nM (bottom) LEN and quantification (right) of VLP diameter along the horizontal axis (technical replicates, n = 56, Wilcoxon Rank-Sum Test, *** P<0.001). **B)** Representative micrographs (left) of CLPs with 0 μM (top) or 500 μM (bottom) LEN and quantification (right) of intact CLP morphology (technical replicates, n = 10, Wilcoxon Rank-Sum Test, *** P<0.001). **C)** Cartoon schematic, model comparison, and RSMD (0.985 Å) between the PR-D25A premature and the PR-WT mature lattices. Zoom-in panel shows that LEN adopts the same conformation between both lattices. **D)** Cartoon schematic, model comparison, and RMSD (0.766 Å) of the PR-D25A premature lattice NTD and the *in vitro* immature lattice NTD. Zoom-in panels show that LEN has rotational freedom at the M3 group. **E)** Viral protein expression in cell lysate (release data in Fig. 2E) measured from 293FT or 293FT-IPPK-KO cells pretreated for 24 hrs with 50 nM LEN at 48 hrs post-transfection with HIV-1 provirus (n=4, Student's T-Test, ** P<0.01).

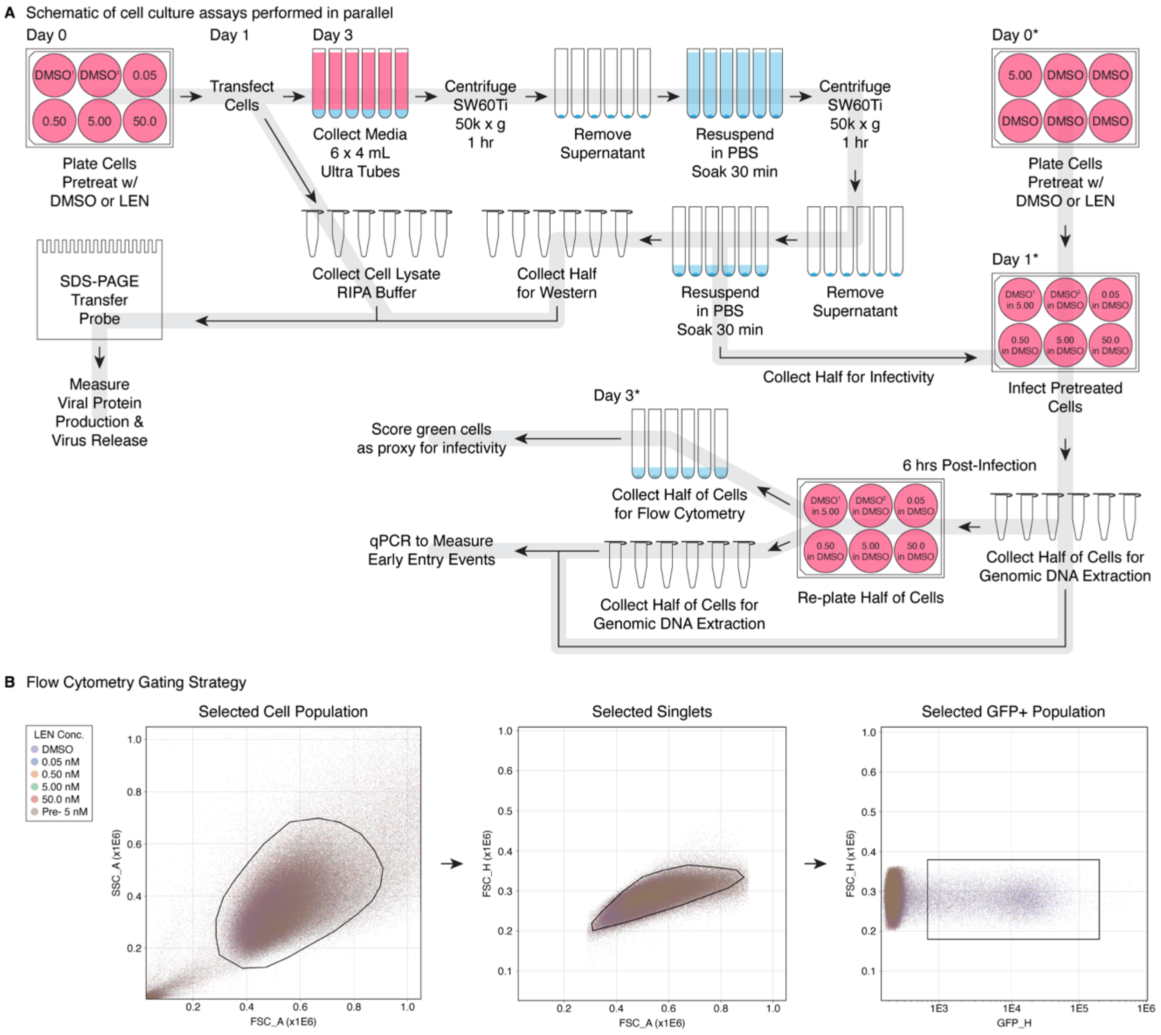


**Figure S9.** *Experimental design of cell culture assays.* **A)** Schematic depicting how samples for viral protein production, viral release, infectivity, and qPCR were collected in parallel from the same experiment with a total of seven biological replicates. **B)** Gating strategy for flow cytometry data from one representative experiment (presented points consist of all treatment groups consolidated on the same plot).

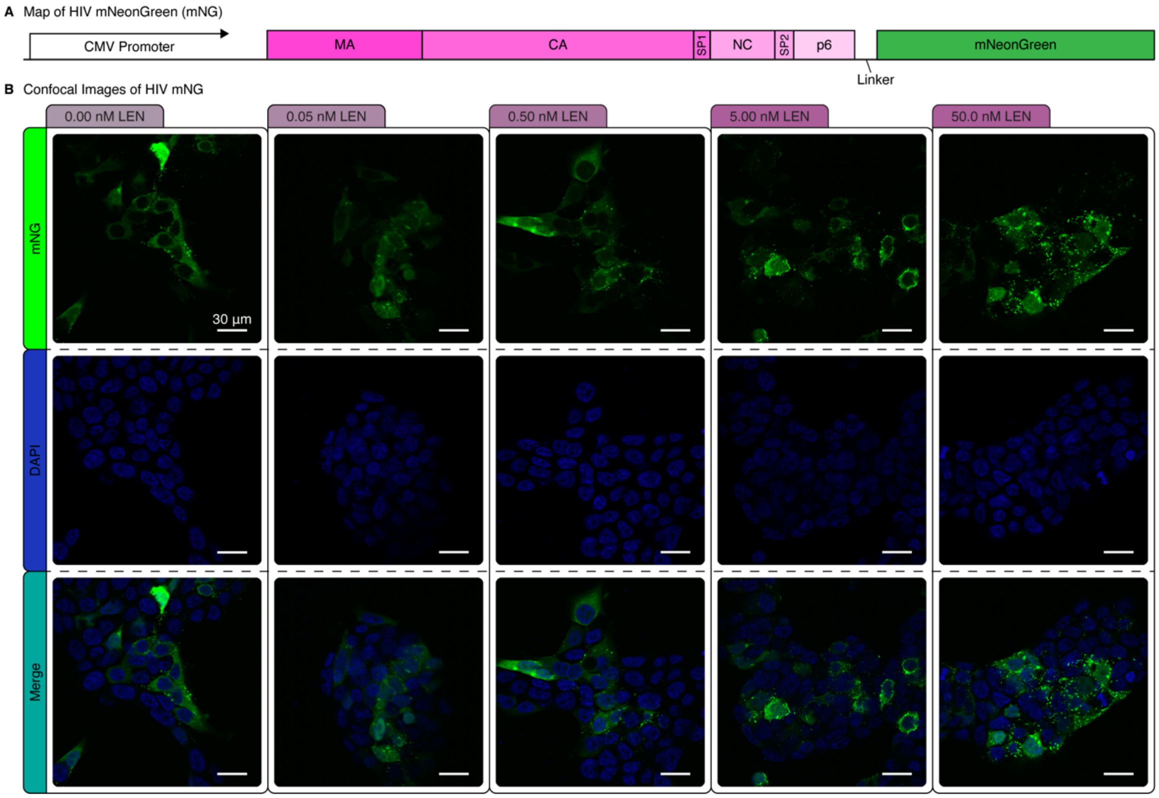


**Figure S10.** *Confocal imaging of cells expressing Gag-mNeonGreen (mNG) fusion protein at 48 hrs post LEN treatment and 24 hrs post transfection.* **A)** Gene map of the HIV-1 Gag-mNG expression construct. **B)** Representative confocal images of cells pretreated with increasing concentrations of LEN (top row: DAPI, middle row: mNG, bottom row: merge).

**Table S1.** Cryo-EM data collection, processing, and model validation statistics.

<table>
<tr><th></th><th></th><th colspan="5">In vitro</th><th colspan="6">In situ</th></tr>
<tr><th></th><th></th><th>CASPNC<br>0 μM LEN</th><th>CASPNC<br>WT 50 μM LEN</th><th>CASPNC<br>WT 50 μM LEN</th><th>CASPNC<br>M66I 50 μM LEN</th><th>CA<br>500 μM LEN</th><th>D25A 5% DMSO<br>(Immature)</th><th>D25A 10 nM LEN<br>(Immature)</th><th>D25A 10 nM LEN<br>(Premature)</th><th>D25A 50 nM LEN<br>(Premature)</th><th>D25A-M66I 50 nM LEN<br>(Immature)</th><th>WT 50 nM LEN<br>(Mature)</th></tr>
<tr><td rowspan="4">Sample Details</td><td>Buffer</td><td>20 mM Tris<br>100 mM NaCl<br>2mM TCEP</td><td>20 mM Tris<br>100 mM NaCl<br>2mM TCEP</td><td>20 mM Tris<br>100 mM NaCl<br>2mM TCEP</td><td>20 mM Tris<br>100 mM NaCl<br>2mM TCEP</td><td>25 mM MES<br>2mM TCEP</td><td>25mM Tris<br>150 mM NaCl</td><td colspan="2">25mM Tris<br>150 mM NaCl</td><td>25mM Tris<br>150 mM NaCl</td><td>25mM Tris<br>150 mM NaCl</td><td>25mM Tris<br>150 mM NaCl</td></tr>
<tr><td>Sample pH</td><td>8</td><td>8</td><td>8</td><td>8</td><td>6.2</td><td>8</td><td colspan="2">8</td><td>8</td><td>8</td><td>8</td></tr>
<tr><td>CASPNC or CA</td><td>50 μM</td><td>50 μM</td><td>50 μM</td><td>50 μM</td><td>500 μM</td><td>NA</td><td colspan="2">NA</td><td>NA</td><td>NA</td><td>NA</td></tr>
<tr><td>LEN</td><td>0 μM</td><td>50 μM</td><td>50 μM</td><td>50 μM</td><td>500 μM</td><td>0 nM</td><td colspan="2">10 nM</td><td>50 nM</td><td>50 nM</td><td>50 nM</td></tr>
<tr><td rowspan="13">Data Collection</td><td>Microscope</td><td>Titan Krios<br>(Thermo)</td><td>Titan Krios<br>(Thermo)</td><td>Talos Arctica<br>(Thermo)</td><td>Talos Arctica<br>(Thermo)</td><td>Titan Krios<br>(Thermo)</td><td>Titan Krios<br>(Thermo)</td><td colspan="2">Titan Krios<br>(Thermo)</td><td>Titan Krios<br>(Thermo)</td><td>Talos Arctica<br>(Thermo)</td><td>Titan Krios<br>(Thermo)</td></tr>
<tr><td>Detector</td><td>K3 (Gatan)</td><td>K3 (Gatan)</td><td>K3 (Gatan)</td><td>K3 (Gatan)</td><td>K3 (Gatan)</td><td>K3 (Gatan)</td><td colspan="2">K3 (Gatan)</td><td>K3 (Gatan)</td><td>K3 (Gatan)</td><td>K3 (Gatan)</td></tr>
<tr><td>Energy Filter</td><td>BioQuantum<br>(Gatan)</td><td>BioContinuum<br>(Gatan)</td><td>BioQuantum<br>(Gatan)</td><td>BioQuantum<br>(Gatan)</td><td>BioContinuum<br>(Gatan)</td><td>BioQuantum<br>(Gatan)</td><td colspan="2">BioQuantum<br>(Gatan)</td><td>BioQuantum<br>(Gatan)</td><td>BioQuantum<br>(Gatan)</td><td>BioContinuum<br>(Gatan)</td></tr>
<tr><td>Energy Filter Slit Width (eV)</td><td>20</td><td>10</td><td>20</td><td>20</td><td>20</td><td>20</td><td colspan="2">20</td><td>20</td><td>20</td><td>20</td></tr>
<tr><td>Nominal Magnification</td><td>105,000 x</td><td>130,000</td><td>63,000 x</td><td>63,000 x</td><td>105,000 x</td><td>105,000 x</td><td colspan="2">105,000 x</td><td>105,000 x</td><td>63,000 x</td><td>105,000 x</td></tr>
<tr><td>Voltage (kV)</td><td>300</td><td>300</td><td>200</td><td>200</td><td>300</td><td>300</td><td colspan="2">300</td><td>300</td><td>200</td><td>300</td></tr>
<tr><td>Total Dose (e-/Å^2)</td><td>50</td><td>40</td><td>50</td><td>50</td><td>50</td><td>50</td><td colspan="2">50</td><td>50</td><td>50</td><td>50</td></tr>
<tr><td>Super-Resolution Mode?</td><td>Yes</td><td>Yes</td><td>Yes</td><td>Yes</td><td>Yes</td><td>Yes</td><td colspan="2">Yes</td><td>Yes</td><td>Yes</td><td>Yes</td></tr>
<tr><td>Acquisition Software</td><td>SerialEM</td><td>SerialEM</td><td>SerialEM</td><td>SerialEM</td><td>SerialEM</td><td>SerialEM</td><td colspan="2">SerialEM</td><td>SerialEM</td><td>SerialEM</td><td>SerialEM</td></tr>
<tr><td>Defocus Range (μm)</td><td>-0.4 to -1.4</td><td>-0.3 to -0.9</td><td>-0.4 to -1.4</td><td>-0.4 to -1.4</td><td>-0.4 to -1.4</td><td>-0.4 to -1.4</td><td colspan="2">-0.4 to -1.4</td><td>-0.4 to -1.4</td><td>-0.4 to -1.4</td><td>-0.4 to -1.4</td></tr>
<tr><td>Pixel Size (Å)</td><td>0.822</td><td>0.6488</td><td>1.31</td><td>1.31</td><td>0.830</td><td>0.822</td><td colspan="2">0.822</td><td>0.822</td><td>1.3</td><td>0.822</td></tr>
<tr><td>Frames Per Movie</td><td>50</td><td>55</td><td>50</td><td>50</td><td>50</td><td>50</td><td colspan="2">50</td><td>50</td><td>50</td><td>50</td></tr>
<tr><td>Number of Movies</td><td>14,686</td><td>18,607</td><td>2,403</td><td>2,558</td><td>7,510</td><td>13,624</td><td colspan="2">7,206</td><td>13,411</td><td>2,690</td><td>12,465</td></tr>
<tr><td rowspan="5">Processing</td><td>Final Number of Particles</td><td>2,468,740</td><td>976,034</td><td>697,457</td><td>422,469</td><td>594,567</td><td>509,305</td><td>63,161</td><td>79,207</td><td>381,307</td><td>146,560</td><td>284,910</td></tr>
<tr><td>Symmetry Imposed</td><td>C6</td><td>C6</td><td>C6</td><td>C6</td><td>C6</td><td>C6</td><td>C6</td><td>C6</td><td>C6</td><td>C6</td><td>C6</td></tr>
<tr><td>B Factor Used for Map Sharpening (Å)</td><td>-61</td><td>-62</td><td>-84</td><td>-80</td><td>-158</td><td>-74</td><td>-61</td><td>-195</td><td>-120</td><td>-98</td><td>-255</td></tr>
<tr><td>Map Resolution at 0.143 FSC (Å)</td><td>1.9</td><td>2.0</td><td>2.6</td><td>2.6</td><td>3.5</td><td>2.3</td><td>2.7</td><td>4.8</td><td>3.1</td><td>3.2</td><td>4.3</td></tr>
<tr><td>EMBD ID</td><td>EMD-75781</td><td>EMD-75782</td><td>EMD-75783</td><td>EMD-75784</td><td>EMD-48430</td><td>EMD-75785</td><td>EMD-75786</td><td>EMD-75787</td><td>EMD-75788</td><td>EMD-75789</td><td>EMD-75790</td></tr>
<tr><td rowspan="11">Atomic Model</td><td>Protein Residues</td><td>639</td><td>856</td><td></td><td>856</td><td>688</td><td></td><td></td><td></td><td>345</td><td></td><td></td></tr>
<tr><td>MolProbability Score</td><td>0.78</td><td>0.69</td><td></td><td>0.68</td><td>1.66</td><td></td><td></td><td></td><td>1.08</td><td></td><td></td></tr>
<tr><td>Clash Score</td><td>0.40</td><td>0.15</td><td></td><td>0.22</td><td>0.6</td><td></td><td></td><td></td><td>0.18</td><td></td><td></td></tr>
<tr><td>Rotamer Outliers (%)</td><td>0.37</td><td>1.11</td><td></td><td>1.11</td><td>4.02</td><td></td><td></td><td></td><td>1.35</td><td></td><td></td></tr>
<tr><td>Ramachandran Favored (%)</td><td>97.29</td><td>97.50</td><td></td><td>97.74</td><td>91.58</td><td></td><td></td><td></td><td>93.77</td><td></td><td></td></tr>
<tr><td>Ramachandran Allowed (%)</td><td>2.55</td><td>2.50</td><td></td><td>2.26</td><td>5.39</td><td></td><td></td><td></td><td>5.93</td><td></td><td></td></tr>
<tr><td>Ramachandran Outliers (%)</td><td>0.16</td><td>0.00</td><td></td><td>0.00</td><td>3.03</td><td></td><td></td><td></td><td>0.30</td><td></td><td></td></tr>
<tr><td>Ramachandran Z-Score</td><td>0.22</td><td>1.69</td><td></td><td>1.74</td><td>-0.45</td><td></td><td></td><td></td><td>2.07</td><td></td><td></td></tr>
<tr><td>RMSD, Bond Length (Å)</td><td>0.019</td><td>0.023</td><td></td><td>0.023</td><td>0.025</td><td></td><td></td><td></td><td>0.021</td><td></td><td></td></tr>
<tr><td>RMSD, Bond Angles (°)</td><td>2.415</td><td>2.570</td><td></td><td>2.518</td><td>3.644</td><td></td><td></td><td></td><td>2.374</td><td></td><td></td></tr>
<tr><td>PDB ID</td><td>11KU</td><td>11KV</td><td></td><td>11KW</td><td>9MNM</td><td></td><td></td><td></td><td>11KX</td><td></td><td></td></tr>
</table>

**Movie S1** (separate file). *Movie demonstrating rotation around αH2 at the $CA_{CTD}$ trimer interface.* Trimer centers and the $CA_{NTD}$ are in dark blue and blue, respectively. The $CA_{CTD}$ and SP are transparent tan and lavender, respectively. IP6 is in maroon and colored by heteroatom.
